# A systematic review and meta-analysis of risk factors for *Leishmania* infection in dogs in Europe

**DOI:** 10.64898/2026.08.03.742436

**Authors:** Erlend I. F. Fossen, Carla Maia, Kristin Aunan

## Abstract

**Background:** Identifying risk factors for *Leishmania* infection in dogs is essential for understanding disease epidemiology and informing control and targeted prevention. This systematic review and meta-analysis examined these risk factors across endemic and non-endemic regions of Europe.

**Methods:** Inclusion criteria included studies that quantified the association between *Leishmania* infection (both asymptomatic infection and canine leishmaniosis) and risk factors among European dogs. Non-peer reviewed grey literature and studies that focus only on prevention were excluded. The search included all records indexed in PubMed up to 17^th^ March 2026. Risk of bias was assessed using the ROBINS-E tool. Random-effects multilevel models were used for meta-analysis, and sensitivity analyses were conducted to assess robustness.

**Results:** Of the 46 studies included, 94% were cross-sectional and 76% assessed seroprevalence. Studies were primarily conducted in Southern Europe (Spain 35%, Portugal 17%, Italy 17%, and Greece 11%). There was considerable evidence of an association between increasing age and infection (OR = 1.15 [95% CI: 1.10, 1.21] per year). Five additional factors showed some evidence of association: male sex (OR = 1.17 [1.01, 1.36]), higher body weight (OR = 1.06 [1.02, 1.09] per kg), outdoor housing (OR = 1.84 [1.26, 2.69]), outdoor sleeping (OR = 2.50 [1.52, 4.09]), and neuter status showing higher odds in neutered dogs (OR = 2.22 [1.37, 3.61]). Other potential risk factors, with weak or no clear evidence, included fur length, coat color, breed, utilization (e.g. pet or hunting), urbanicity, care setting, clinical signs, cohabitation, co-infection, sand fly density, land cover, home features, owner perception and knowledge of the disease, travel history and origin of dog, and area-level socioeconomic deprivation. Over 75% of analytical units were assessed as having a high risk of bias, with the remaining assessed as moderate risk. Inadequate handling of confounders was the primary source of bias risk.

**Conclusions:** The results highlight key risk factors to consider in future research and for improving prevention and disease control, with most identified factors likely acting as proxies of exposure to sand fly bites. Future studies should ensure adequate control for confounding, while causal inference would be strengthened by conducting longitudinal cohort studies.

## Background

Canine leishmaniosis is an infectious vector-borne disease affecting dogs worldwide, with at least 2.5 million infected dogs in Southwestern Europe[1]. The disease is caused by protozoans from the genus *Leishmania*, and transmitted through bites from infected females of the hematophagous sand fly (Diptera, Phlebotominae)[2]. Zoonotic visceral leishmaniosis is primarily caused by *L. infantum*, with humans as incidental hosts and dogs as the primary reservoir[3]. Clinical signs in dogs, while not always present, vary considerably, but often include dermatological lesions, lymphadenomegaly, weight loss, ocular abnormalities and renal disease[3]. Given the close relationship between humans and dogs, canine leishmaniosis is of importance for both veterinary and public health, making identification of risk factors in dogs essential for disease control and the development of targeted prevention.

Canine leishmaniosis is endemic in Southern Europe[4], but increasing number of cases are being reported in previously non-endemic areas. This reflects both the emergence of autochthonous transmission, driven partly by climatic changes that favor sand fly vectors, and the introduction of infection through the movement of dogs and increased human mobility within the European Union[5]. With further expansion expected under ongoing climate changes[6, 7], these trends highlight the importance of identifying risk factors for infection in dogs across Europe, encompassing both traditionally endemic regions and areas where the disease is emerging.

Previous reviews and meta-analyses of risk factors have partly focused on the Americas, particularly Brazil, where transmission dynamics, vector species and ecological conditions differ considerably from those in Europe[8, 9]. Other reviews have focused on North African countries[10, 11] or on Iran[12], representing settings with different socio-economic conditions, patterns of dog ownership, access to veterinary care, and disease surveillance infrastructure compared to the European context[10, 12, 13]. In addition, reviews with a global[14] or Mediterranean scope[15] have focused on disease prevalence and its variation across canine populations, rather than formally quantifying the association between risk factors and infection. Consequently, systematic synthesis and quantitative meta-analytic assessment of risk factors specific to the European context remain lacking, despite considerable variation in disease distribution across Europe, including endemic areas and regions with sporadic or emerging transmission.

The aim of this study was to conduct a systematic review and meta-analysis of risk factors for *Leishmania infantum* infection in dogs in Europe, providing quantitative assessment across both endemic and non-endemic settings. By doing so, we provide the first synthesis of this kind focused specifically on Europe, contributing to improved understanding of disease epidemiology and informing targeted prevention and control strategies.

## Methods

### Study design

This systematic review and meta-analysis was performed following the guidelines of the 2020 Preferred Reporting Items for Systematic Reviews and Meta-Analyses (PRISMA[16]). The checklist is provided in Additional file 1. The literature search aimed to identify all relevant studies that mention *Leishmania* infection in dogs (including both asymptomatic infection and canine leishmaniosis), risk factors for infection, and Europe or a European country where the disease is reported. The meta-analysis and narrative review aimed to synthesize which risk factors are associated with *Leishmania* infection in dogs, and their overall effect sizes. A single author screened the literature, collected data and assessed risk of bias for all relevant studies. This review was not registered beforehand. A formal protocol was not published.

### Literature search

The literature search was conducted in PubMed, which provides extensive coverage of biomedical and veterinary research. Two searches were conducted by one researcher, first on 20^th^ February 2025, and an additional search on 17^th^ March 2026 to include any newly added publications. Overall, the search included all records indexed in PubMed up to 17^th^ March 2026. The search string explicitly listed all European countries where the disease may have been reported, based on disease distributions found in four relevant publications[4, 13, 17, 18]. The following search string was used in PubMed:

((canine OR dog OR dogs) AND (leishmani*) AND (spain OR portugal OR iberia* OR mediterra* OR europe OR france OR italy OR germany OR greece OR switzerland OR UK OR “united kingdom” OR belgium OR netherlands OR albania OR bulgaria OR serbia OR bosnia* OR montenegro OR cyprus OR turkey OR croatia OR sweden OR denmark OR norway OR finland OR hungary OR romania OR macedonia OR slovenia OR malta OR kosovo OR austria OR czechia OR slovakia OR ukraine OR moldova OR “san marino”) AND (risk* OR vulnerab* OR “associated factor*” OR “epidemiological stud*” OR “epidemiological factor*”))

### Study selection and eligibility criteria

All studies were screened by one author in two stages. In the 1^st^ stage the title, abstract and keywords were screened and excluded based on the following exclusion criteria:

1. Study not about *Leishmania* infection in dogs
2. Study does not consider risk factors for infection
3. Study does not include data on dogs from Europe
4. Study is grey literature (e.g. report from symposium) and not published in a peer-reviewed journal
5. Study only contains risk factors related to treatment, preventives or genetic traits

The remaining studies after the first stage of screening underwent full-article screening in the 2^nd^ stage. In addition to the previously stated exclusion criteria, the following exclusion criteria were used:

6. Effect sizes for risk factors were not obtainable from the study (i.e. neither reported nor derivable from available data).
7. Study is a review without any new effect size estimates
8. Study deemed to have too high risk of bias after bias assessment

Among the selected studies, all risk factors that relate to the effectiveness of treatment or preventive measures (e.g. vaccination, use of collars or repellents), genetic traits, or that is only relevant in that specific context (e.g. locations such as south vs north, altitude, and climate variables), were excluded from downstream analyses. Treatment, preventive measures and genetic traits were considered out of scope for this review (see e.g. [19] for a review on vaccination efficacy). Context-specific variables were excluded because they are not directly comparable across studies.

Some studies were subdivided into multiple “sub studies”, defined as distinct groups or strata within a study for which results were reported separately (e.g. based on population, study design, diagnostic method, outcome definition, or time period). Studies without such subdivisions were each treated as a single sub study.

### Data collection from eligible studies

#### Collecting study characteristics

For each sub-study, we collected publication information (e.g. authors, year), country and geographical region, study design (e.g. cross-sectional), diagnostic test (e.g. indirect immunofluorescence antibody test (IFAT) or polymerase chain reaction (PCR)), disease outcome (e.g. seropositivity or current infection), sample sizes (total tested and number of positive dogs), and whether the study controlled for confounders. Additionally, we collected the names and categories of all risk factors that were reported in each study, which were subsequently harmonized and analyzed in meta-analyses.

#### Risk of bias assessment

Risk of bias was assessed using the Risk Of Bias In Non-randomized Studies of Exposures (ROBINS-E) tool[20]. In this review, candidate risk factors are considered as exposures. These assessments were done on the level of analytical units within each sub-study, where analytical units were defined as individual risk-factor estimates (unadjusted or adjusted). Sub-studies reporting only one type of analysis contributed a single unit, while those reporting both were split into two analytical units, with separate assessments reflecting differences in risk of confounding.

Risk of bias was assessed using modified versions of the seven domains of the ROBINS-E tool (“confounding”, “measurement of risk factors”, “selection bias”, “post exposure interventions”, “missing data”, “measurement of outcome”, and “reporting bias”) and one additional domain (“other bias”) to include any additional source of bias not captured by the tool. Under the “confounding” domain, analytical units based on univariable (unadjusted) analyses were considered at high risk of bias for not addressing confounding. Analytical units that presented multivariable (adjusted) analyses, but excluded possible confounders based solely on their non-significance in univariable analysis or through stepwise variable selection, were considered at medium risk of bias in the “confounding” domain since such approaches are known for giving inflated effect estimates and result in invalid confidence intervals and p-values[21, 22]. Analytical units that included possible confounders and reported adjusted estimates without excluding non-significant factors were considered at low risk of bias under the “confounding” domain. Because IFAT is considered the reference method for *Leishmania* serology in dogs[23], under the “measurement of outcome” domain, for studies reporting seroprevalence, only studies using IFAT for diagnosis were considered at low risk of bias, while the remaining serological alternatives were considered at medium risk of bias. The overall risk of bias was considered the same as the highest risk in any domain, while allowing for an additive judgment of very high risk of bias when multiple domains were rated as high risk, in accordance with ROBINS-E guidelines. Based on this principle, analytical units with high risk of bias in four or more domains were excluded from quantitative synthesis.

#### Harmonization of risk factors

To enable comparison across studies, reported risk factors were harmonized by grouping conceptually similar variables and standardizing their definitions. Overlapping variables were aligned under common labels using regrouping or dichotomizing when possible and relevant. The most frequent definition was used in cases where several similar but not harmonizable definitions were available. Alternative definitions (coding) of risk factors were studied in sensitivity analyses. Risk factors, as well as individual definitions of risk factors, with fewer than three comparable effect estimates did not undergo formal meta-analysis but were included in the narrative synthesis of the literature. Below follows specifics on how risk factors were harmonized.

##### Age

Age was reported very heterogeneously across studies, including both continuous measures and multiple different categorical definitions (e.g. “young” vs “adult”, <2 years vs >2 years, or grouped into 2-year age groups). The most frequent reporting formats allowed derivation of age as a continuous variable (per-year increase), which was used in the main analysis to maximize the number of sub-studies contributing estimates. Three alternative categorical codings were explored in sensitivity analyses: <1 year vs ≥1 year; ≤7 years vs >7 years; <3 years vs 3-4 years/3-5 years/3-6 years (hereby referred to as <3 vs 3-6 years).

##### Sex

Sex was reported as female vs male and required no further harmonization.

##### Weight

Weight was reported heterogeneously across studies, including both continuous measures and several categorical definitions (e.g. “small” vs “large”, <25 kg vs >25 kg, or grouped into 10-kg weight groups). The most frequent reporting formats allowed derivation of weight as a continuous variable (per-kg increase), which was used in the main analysis to maximize the number of sub-studies contributing estimates. Four alternative categorical codings were explored in sensitivity analyses: <25 kg vs >25 kg; “small” vs “large”; “small” vs “medium”; “medium” vs “large”. Note that the latter three only include sub-studies that reported all three weight categories (small, medium, large), excluding sub-studies reporting only “small” vs “large”, as these do not account for the medium category and are therefore not directly comparable.

##### General housing

General housing, here defined as the dogs primary housing type, was reported along a gradient from exclusively indoors to exclusively outdoors, with varying categorization (e.g. “mainly indoor” vs “mainly outdoor”, or “indoor” vs “mixed” vs “outdoor”). The most frequent definition was “indoors” vs “outdoors”, without a “mixed” category. For harmonization, the main analyses used indoors vs outdoors as the comparison, where definitions using “mostly” or “mainly” indoors/outdoors were classified as indoors or outdoors, respectively. Two alterative codings were explored in sensitivity analyses: “partly outdoors” (mixed) vs “outdoors”; “indoors” vs “partly outdoors”/“outdoors”.

##### Sleeping location

Sleeping location was reported as indoors vs outdoors and required no further harmonization.

##### Fur length

Fur length, or coat/hair length, was reported along a gradient from short to long, with varying categorization (e.g. short vs long, short vs medium vs long, or short vs medium/long). The most frequent definition was short vs medium/long, which was used in the main analysis, as it was also harmonizable with studies reporting medium and long fur as separate categories. As alternative coding, a binary categorization of short vs long was used for studies that did not report a medium category. This was analyzed separately, as the absence of a medium category precludes direct comparison with definitions including three levels.

##### Neuter status

Neuter status was reported as entire (intact) vs neutered and required no further harmonization.

##### Breed

Breed was reported very heterogeneously across studies, including studies reporting broad categories (e.g. mixed, purebred, mongrel), studies reporting origin (e.g. indigenous to a specific country, autochthonous or exotic) and studies reporting individual breeds (e.g. German Shepards, terriers). For harmonization, named breeds were classified as pure, while mongrel and mixed breeds were treated as separate categories only when explicitly distinguished in the studies. The most frequent harmonized comparisons were pure vs mixed/mongrel, followed by mongrel vs combined pure/mixed, both of which were used in the main analyses. Three alternative codings were explored in sensitivity analyses, using studies that reported all three broad categories (pure, mixed, mongrel): mixed vs pure; mongrel vs pure; mixed vs mongrel.

##### Utilization

Utilization, or dog function, was reported heterogeneously across studies, including categories such as working dogs, pets, sheepdogs, hunting dogs, guard dogs and breeding dogs. The most frequent harmonized comparisons were between hunting dogs vs non-hunting dogs, and between pets (including companion dogs) and non-pets. As such, these comparisons were the ones used in the analyses.

##### Urbanicity

Urbanicity, the degree of urbanization, was reported heterogeneously along a gradient from fully rural to fully urban, with varying categorization (e.g. broadly as rural vs urban, and more narrowly as high-density urban vs isolated ranches with several categories in between). The most frequent harmonized comparison was rural vs periurban/urban, which was used in the main analysis, as it was also harmonizable with studies reporting periurban, urban, and other degrees of urbanicity as separate categories. Two alternative codings were explored in sensitivity analyses: periurban vs rural; a binary categorization of urban vs rural, for studies that did not report a medium category (periurban). The latter was analyzed separately, as the absence of a medium category limits direct comparability with definitions that included intermediate levels of urbanicity.

##### Care setting

Care setting, defined as the dog’s living context (including ownership and housing), was reported heterogeneously, including categories such as living at home, being owned, living in a shelter or kennel, being stray, living on a farm or being domestic. The most frequent comparison was between dogs living at home and stray dogs, which was therefore used in the main analysis. One alternative coding was explored in sensitivity analyses, namely dogs living at home vs dogs in other care settings (combined category).

##### Living with other dogs

Cohabitation with other dogs in general was reported either as binary (yes vs no) or as the number of dogs in the household. For harmonization, dogs in single-dog households were classified as not cohabiting, while dogs in households with >1 dog were classified as cohabiting. A binary categorization (yes vs no) was used in the analysis. Living with seropositive dogs was considered a distinct risk factor and was therefore not included in the broader category of cohabitation with dogs.

##### Clinical signs

Clinical signs, here used broadly as an overall clinical status classification, were reported heterogeneously across studies, with comparisons such as symptomatic vs asymptomatic, healthy vs sick, suspect vs non-suspect, and clinical signs of vector-borne disease (yes vs no). For harmonization, any indication of impaired clinical status was classified as presence of clinical signs and analyzed as a binary variable (yes vs no).

##### Other factors

All other risk factors had fewer than three comparable effect estimates and were therefore not included in the harmonization process. These were instead included in the narrative synthesis of the review.

#### Collecting effect sizes per risk factor

Odds ratios (OR) with 95% confidence intervals were the effect sizes of interest for the meta-analysis. We extracted OR directly when reported or calculated them from other reported data. This included calculating OR from log OR with standard errors, and from crude counts (or percentages) of infected/seropositive dogs by risk factor category along with total number of dogs tested per category. Crude counts or percentages were only collected for the purpose of calculating OR of unadjusted estimates, whereas adjusted estimates were reported by studies only as OR and were extracted as such. For risk factors or definitions of risk factors that were not harmonized, the reported direction and statistical significance were collected.

### Statistical analyses

All statistical analyses were performed in R Statistical Software[24] (v4.6.0).

#### Standardization of effect sizes to odds ratios

When OR were not directly reported, they were calculated from available data. Log ORs and their 95% confidence intervals (CI) were exponentiated. 2x2 contingency tables were reconstructed to calculate OR and 95% CI when only crude counts or percentages were reported. When studies reported only proportions, the number of events was approximated based on the reported percentages and sample size. A continuity correction of 0.5 was applied for 2x2 tables with zero cells. Where necessary, effect estimates were inverted to ensure consistent reference category across studies.

For age and weight, categorical data were transformed into continuous variables. ORs were first calculated (if not already present) for each reported category comparison, then exposure levels were assigned using category midpoints based on reported lower and upper bounds. For example, if a study compared 0-1 years to 2-3 years, the midpoints would be 0.5 and 2.5 years for each of these. For open-ended categories (e.g. >7 years), upper bounds were approximated by adding the largest observed interval width from the same study. This approach assumes that the width of the highest category is comparable to the adjacent intervals and reflects commonly used approximations when handling open-ended exposure categories in dose-response analyses[25]. Study-specific OR estimates per 1-unit increase (year or kg) were then calculated using fixed-effect dose-response meta-analysis with the “dosresmeta” package[26] in R. This was only applied to studies reporting at least two exposure contrasts, allowing estimation of dose-response relationships. Studies reporting only single comparisons (e.g. <2 vs >2 years) were not included in this transformation. Ultimately, these study-specific estimates were included in the meta-analyses together with other reported continuous OR estimates.

#### Meta-analysis

Meta-analyses were performed in R using the “metafor” package[27]. Primary analyses were defined as random-effects multilevel models based on adjusted effect estimates for the main harmonized definitions, where available. Primary analyses were only conducted for risk factors that had adjusted estimates from a minimum of three sub-studies. Secondary analyses were defined as the corresponding random-effects multilevel models based on unadjusted effect estimates for the same definitions and are reported distinctly throughout. Additional analyses, including those based on alternative codings of risk factors, are reported under sensitivity analyses.

Given that multiple effect estimates (sub-studies) could originate from the same study, a multilevel modelling approach was applied, with sub-studies nested within studies, using the “rma.mv()” function. Models were fitted using restricted maximum likelihood (REML). This approach uses a multivariate weighting structure that accounts for within-study dependence, such that studies contributing multiple sub-studies do not disproportionately influence the pooled estimates. Statistical heterogeneity was quantified using I^2^, decomposed into between-study and within-study variance components, where the latter reflects variability between sub-studies from the same study. Total I^2^ reflects the sum of both components. Results were visualized using forest plots.

#### Publication bias

Assessment of potential reporting bias (publication bias) was conducted for analyses including at least 10 studies. Publication bias was evaluated at the effect-estimate level, as multiple effect estimates could originate from the same study. Visual inspection of funnel plots was performed, complemented by rank correlation tests. As Egger’s regression test of funnel plot asymmetry is not directly applicable to multilevel models, it was instead applied to corresponding random-effects models without the multilevel structure. Funnel plots from these models were compared with those from the multilevel models and found to be similar. All publication bias analyses should be interpreted with caution, given the small sample sizes and dependence structure of the data.

#### Sensitivity-analyses

To test the robustness of the primary and secondary analyses, several sensitivity analyses were conducted, as described below.

##### Risk of bias

For primary analyses, the risk of bias per sub-study was included as a covariate in corresponding meta-regression models to test if the pooled effect size depended on the risk of bias. This was not done for secondary analyses, as these were all classified as having high risk of bias.

##### Fixed-effects models

Fixed-effects models of corresponding random-effects models were tested to see if pooled estimates vary considerably. Large differences would suggest considerable heterogeneity stemming from between-study and/or within-study (i.e. between sub-study) variance.

##### Mixed-estimates analysis

To partly control for potential reporting bias, where studies do not report non-significant effect sizes in multivariable models, a mixed-estimates analysis was conducted for risk factors with available primary analyses. The mixed-estimates analysis involved using the same adjusted estimates as in the primary analysis but supplementing it with unadjusted estimates from studies where the risk factor was excluded from multivariable models, either due to non-significance in univariable analyses or removed through variable selection. This approach assumes that the effect size of these risk factors would remain similar in a multivariable context and will typically give more conservative pooled estimates.

##### Alternative codings

Alternative codings (definitions) of risk factors were analyzed in separate models to see the influence of decisions regarding definitions of risk factors.

##### Non-harmonized studies

For any given risk factor, a narrative synthesis was conducted of estimates from studies that were not harmonized. Additionally, for each risk factor, we noted how many studies excluded presenting adjusted estimates due to the estimate not reaching statistical significance.

#### Certainty assessment

The robustness of evidence was evaluated by examining confidence interval widths, between- and within-study heterogeneity (I^2^), consistency across sensitivity analyses, risk of bias and publication bias, and the narrative synthesis of studies not included in meta-analyses. Given the observational nature and heterogeneity of the included studies, even the most robust findings reflect moderate certainty at best. We used “considerable evidence” for the most robust findings, followed by “some evidence”, “weak evidence” and “no clear evidence”.

## Results

### Study selection

A total of 434 articles were identified through the PubMed search, two of which were removed as duplicate records. After the first stage of screening, 138 reports were retrieved and underwent full-text screening for eligibility. One seemingly eligible study[28] did not report clear effect sizes and was therefore considered to have a too high risk of bias, and was consequently excluded from downstream analyses. Of the remaining reports, a total of 46 studies were eligible and included in the narrative literature synthesis, and 40 of these studies were included in the formal meta-analyses after harmonizing the risk factors. The PRISMA flowchart (Fig. 1) summarizes the search and selection process. Within the 46 studies, a total of 68 sub-studies were identified and included in the literature review.

**Fig. 1.**
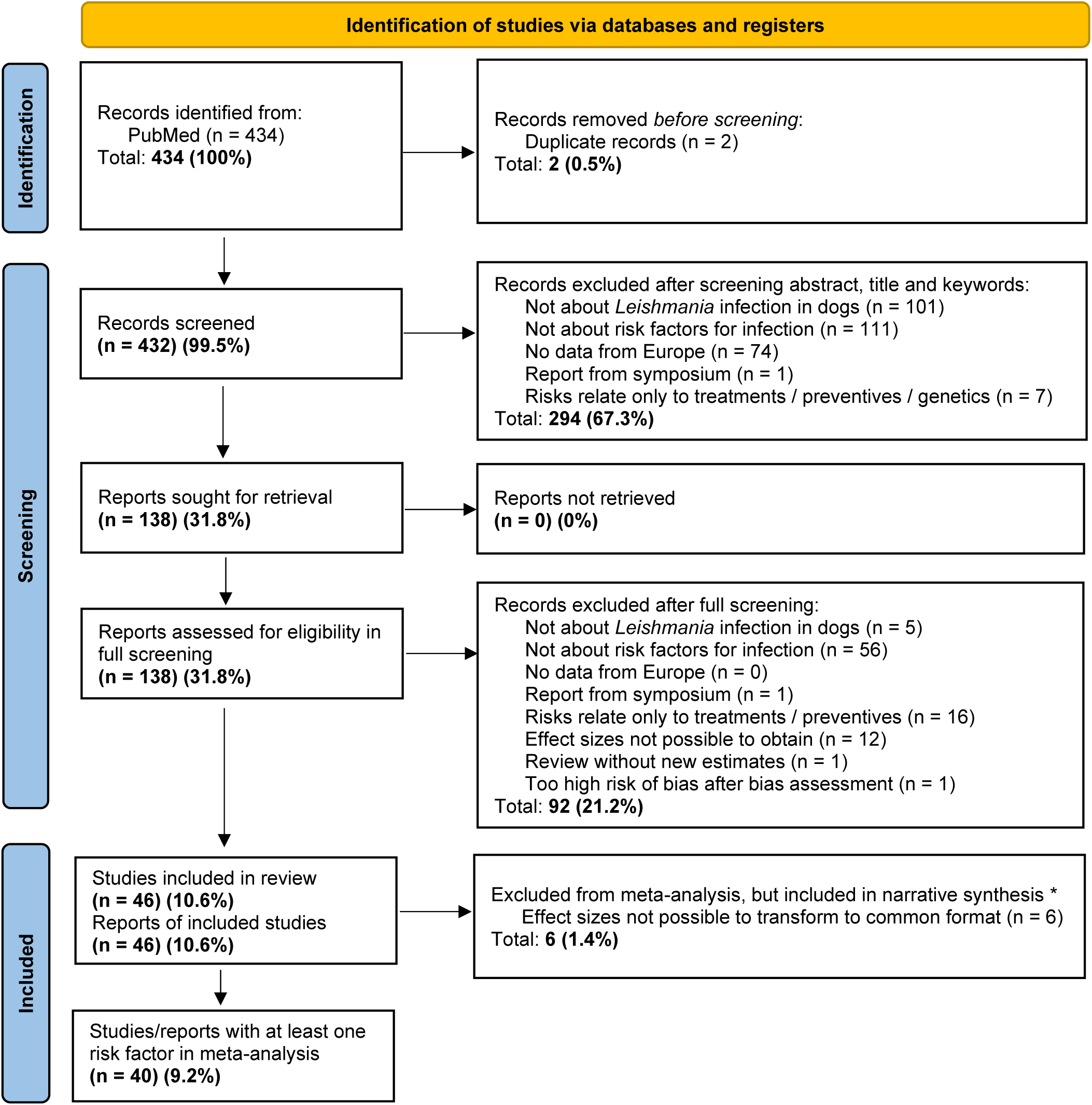
PRISMA flowchart of the study selection process showing inclusion and exclusion of studies. * All studies that passed eligibility criteria (N = 46) are included in the narrative part of the review, even if excluded from the meta-analysis.

### Study characteristics

The included studies (n = 46) were published from year 1995 to 2026, with 59% of the studies being published after 2014 (Additional file 2: Fig. S1). Studies were primarily from Southern Europe with the highest number of studies coming from Spain, followed by Italy, Portugal and then Greece (Table 1). A majority of studies used a cross-sectional design and considered seropositivity as their disease outcome. IFAT was the most used diagnostic test, and approximately half of the studies reported adjusted multivariable analyses of risk factors. Each included study consisted of one or more sub studies. These sub studies displayed similar characteristics to the overall study set but were, on average, published two years later, indicating that newer studies more often reported multiple analyses within a single publication (Table 1). Characteristics of each included study[29–74] is given in Additional file 3: Dataset S1.

**Table 1.** Overview of study characteristics. Sub-studies are nested within studies, where each study (publication) contains at least one sub-study. *Studies assessing current infection using PCR.

|  | STUDIES (N= 46) | SUB-STUDIES (N= 68) |
| --- | --- | --- |
| Publication year |  |  |
| Mean (SD) | 2015.1 (7.2) | 2016.8 (7.0) |
| Median [Min, Max] | 2016.9 [1995, 2026] | 2019.0 [1995, 2026] |
| Country |  |  |
| Albania | 1 (2.2%) | 1 (1.5%) |
| Bosnia and Herzegovina | 2 (4.3%) | 3 (4.4%) |
| France | 1 (2.2%) | 1 (1.5%) |
| Georgia | 1 (2.2%) | 1 (1.5%) |
| Germany | 1 (2.2%) | 2 (2.9%) |
| Greece | 5 (10.9%) | 5 (7.4%) |
| Italy | 8 (17.4%) | 14 (20.6%) |
| Portugal | 8 (17.4%) | 14 (20.6%) |
| Romania | 1 (2.2%) | 1 (1.5%) |
| Spain | 16 (34.8%) | 18 (26.5%) |
| United Kingdom | 2 (4.3%) | 8 (11.8%) |
| Study design |  |  |
| Case-control | 1 (2.2%) | 7 (10.3%) |
| Cross-sectional | 43 (93.5%) | 60 (88.2%) |
| Longitudinal | 0 (0%) | 1 (1.5%) |
| Cross-sectional + Case-control | 1 (2.2%) | 0 (0%) |
| Cross-sectional + Longitudinal | 1 (2.2%) | 0 (0%) |
| Diagnostic tests |  |  |
| Direct agglutination test (DAT) | 4 (8.7%) | 4 (5.9%) |
| Enzyme-linked immunosorbent assay (ELISA) | 6 (13.0%) | 8 (11.8%) |
| IFAT | 20 (43.5%) | 32 (47.1%) |
| Lateral flow test | 4 (8.7%) | 4 (5.9%) |
| Multi-serology | 3 (6.5%) | 3 (4.4%) |
| PCR | 3 (6.5%) | 7 (10.3%) |
| PCR + Serology | 6 (13.0%) | 10 (14.7%) |
| Disease outcome |  |  |
| Current infection* | 4 (8.7%) | 8 (11.8%) |
| Seropositivity | 35 (76.1%) | 50 (73.5%) |
| Seropositivity + Current infection* | 3 (6.5%) | 0 (0%) |
| Seropositivity + Other | 1 (2.2%) | 0 (0%) |
| Other | 3 (6.5%) | 10 (14.7%) |
| Adjusted analysis present? |  |  |
| No | 22 (47.8%) | 36 (52.9%) |
| Yes | 24 (52.2%) | 32 (47.1%) |

### Risk of bias assessment

Based on the ROBINS-E tool, >75% of analytical units within sub-studies were assessed to have high risk of bias, with the remaining rated as having moderate risk (Fig. 2). This was primarily driven by analytical units having high risk of bias related to confounding.

**Fig. 2.**
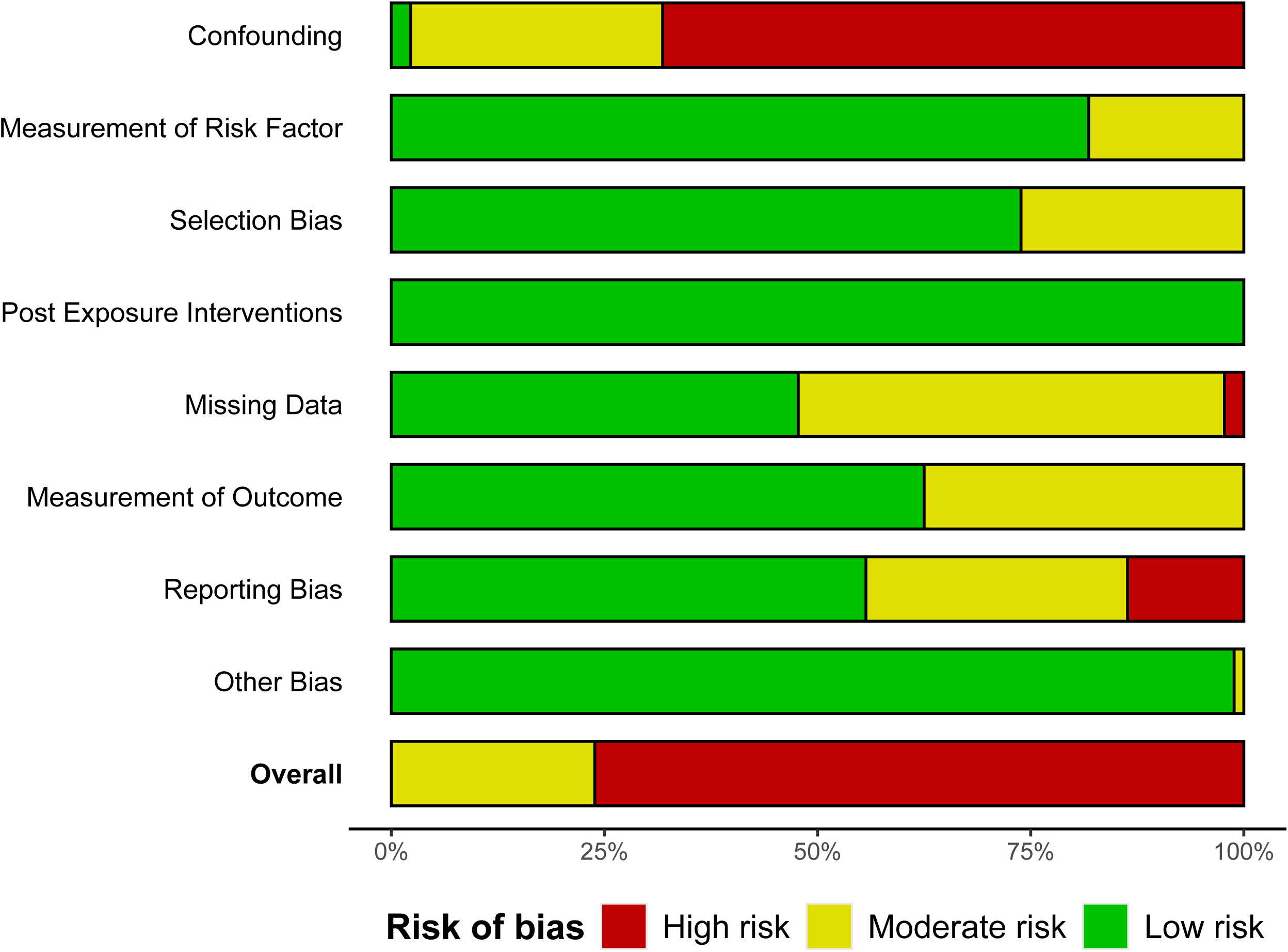
Summary of ROBINS-E risk of bias assessment. Note that risk of bias is summarized per analytic unit within sub studies, with separate assessments for unadjusted and adjusted analyses, and not per study or sub-study.

Analytical units based on univariable analyses were consistently rated as high risk of bias in the confounding domain, while most multivariable analyses were rated as moderate risk of bias because of incomplete covariate adjustments. See Additional file 2: Fig. S2 for a full risk of bias assessment of each study.

### Risk factors and their association with *Leishmania* infection in dogs

Results are described for each risk factor, with the primary analyses (adjusted estimates) being presented in Table 2 and the secondary analyses (unadjusted estimates) presented in Table 3. Effect sizes from each individual sub-study are found in Additional file 4: Dataset S2. Detailed sensitivity analyses are given in Additional file 2: Appendix. For most risk factors, unadjusted estimates were available from more studies than adjusted estimates, as not all studies reported adjusted estimates.

**Table 2.** Adjusted risk factors for *Leishmania* infection in dogs. Pooled odds ratios (OR) from random-effects and fixed-effects meta-analyses evaluating associations between risk factors and *Leishmania* infection based on *adjusted effect estimates only*, with corresponding 95% confidence intervals (CI), number of studies/estimates (k), and measures of total heterogeneity (I^2^). For each comparison, the first category listed represents the reference group. Asterisks (*) indicate statistically significant associations (p<0.05).

| Risk factor | Comparison | k (studies /<br>sub-studies) | Random OR<br>[95% CI] | Global<br>$I^2$ (%) | Fixed OR [95%<br>CI] |
| --- | --- | --- | --- | --- | --- |
| Age of dog | Per 1-year increase | 11 / 12 | 1.15* [1.10, 1.21] | 77.8 | 1.08* [1.06, 1.09] |
| Sex of dog | Female vs male | 4 / 5 | 1.17* [1.01, 1.36] | 52.1 | 1.09* [1.05, 1.13] |
| Weight of dog | Per 1-kg increase | 3 / 3 | 1.06* [1.02, 1.09] | 59.1 | 1.04* [1.03, 1.06] |
| General housing | Indoors vs outdoors | 4 / 4 | 1.84* [1.26, 2.69] | 47.2 | 1.61* [1.29, 2.01] |
| Sleeping location | Indoors vs outdoors | 3 / 3 | 2.50* [1.52, 4.09] | 23.7 | 2.45* [1.62, 3.69] |
| Fur length | Short vs long | 2 / 3 | 0.60 [0.28, 1.30] | 94.5 | 0.84* [0.77, 0.92] |
| Neuter status | Entire vs neutered | 2 / 6 | 2.22* [1.37, 3.61] | 91.2 | 2.24* [1.94, 2.58] |

**Table 3.** Unadjusted risk factors for *Leishmania* infection in dogs. Pooled odds ratios (OR) from random-effects and fixed-effects meta-analyses evaluating associations between risk factors and *Leishmania* infection based on *unadjusted effect estimates only*, with corresponding 95% confidence intervals (CI), number of studies/estimates (k), and measures of total heterogeneity (I^2^). For each comparison, the first category listed represents the reference group. Asterisks (*) indicate statistically significant associations (p<0.05).

| Risk factor | Comparison | k (studies / sub-studies) | Random OR [95% CI] | Global $I^2$ (%) | Fixed OR [95% CI] |
| --- | --- | --- | --- | --- | --- |
| Age of dog | Per 1-year increase | 27 / 30 | 1.06* [1.02, 1.10] | 69.7 | 1.05* [1.05, 1.06] |
| | < 1 vs ≥ 1 years | 13 / 14 | 2.10* [1.13, 3.89] | 46.4 | 2.15* [1.74, 2.65] |
| | ≤ 7 vs > 7 years | 11 / 13 | 1.27 [0.95, 1.70] | 60.9 | 0.91 [0.82, 1.00] |
| Sex of dog | Female vs male | 31 / 39 | 1.20* [1.12, 1.28] | 11.7 | 1.13* [1.10, 1.17] |
| Weight of dog | Per 1-kg increase | 9 / 9 | 1.02* [1.01, 1.04] | 67.0 | 1.02* [1.01, 1.02] |
|  | < 25 vs > 25 kg | 5 / 5 | 1.42 [0.85, 2.36] | 73.7 | 1.07 [0.88, 1.30] |
|  | Small vs large | 6 / 7 | 2.01* [1.23, 3.21] | 55.0 | 1.11* [1.03, 1.20] |
| General housing | Indoors vs outdoors | 9 / 9 | 2.16* [1.20, 3.88] | 57.0 | 1.49* [1.35, 1.63] |
| Sleeping location | Indoors vs outdoors | 7 / 7 | 2.12* [1.42, 3.17] | 30.2 | 1.31* [1.17, 1.46] |
| Fur length | Short vs medium/long | 8 / 14 | 0.84* [0.71, 0.98] | 9.8 | 0.85* [0.76, 0.95] |
| Neuter status | Entire vs neutered | 3 / 3 | 2.62* [2.16, 3.17] | 13.9 | 2.70* [2.36, 3.54] |
| Breed | Pure vs mixed/mongrel | 17 / 25 | 1.05 [0.78, 1.42] | 76.5 | 1.14* [1.10, 1.19] |
|  | Mongrel vs pure/mixed | 6 / 12 | 1.35* [1.07, 1.71] | 14.4 | 1.37* [1.16, 1.61] |
| Utilization | Hunting vs other | 9 / 9 | 0.68* [0.46, 1.00] | 73.3 | 0.77* [0.68, 0.87] |
|  | Pets vs other | 10 / 10 | 1.50* [1.13, 2.00] | 39.1 | 1.53* [1.33, 1.76] |
| Urbanicity | Rural vs periurban/urban | 6 / 6 | 0.70* [0.51, 0.97] | 16.4 | 0.82* [0.70, 0.96] |
| Care setting | Home vs stray | 4 / 5 | 0.77* [0.61, 0.97] | 0.0 | 0.77* [0.61, 0.97] |
| Living with other dogs | Yes vs no | 5 / 5 | 0.79* [0.69, 0.90] | 0.0 | 0.79* [0.69, 0.90] |
| Clinical signs | Yes vs no | 9 / 15 | 0.27* [0.16, 0.44] | 71.0 | 0.22* [0.19, 0.26] |

#### Age of dog

The final analytic dataset contained estimates of the association between age and *Leishmania* infection from a total of 29 studies (33 sub-studies). The primary analysis, restricted to adjusted estimates (n = 12 sub-studies), showed a significant association, with the odds of infection increasing by 15% per one year increase in age (p <0.001, Table 2, Fig. 3). There was evidence of heterogeneity (I^2^ = 77.8%, Q = 243.3, df = 11, p < 0.001), with all heterogeneity attributable to between-study variance.

**Fig. 3.**
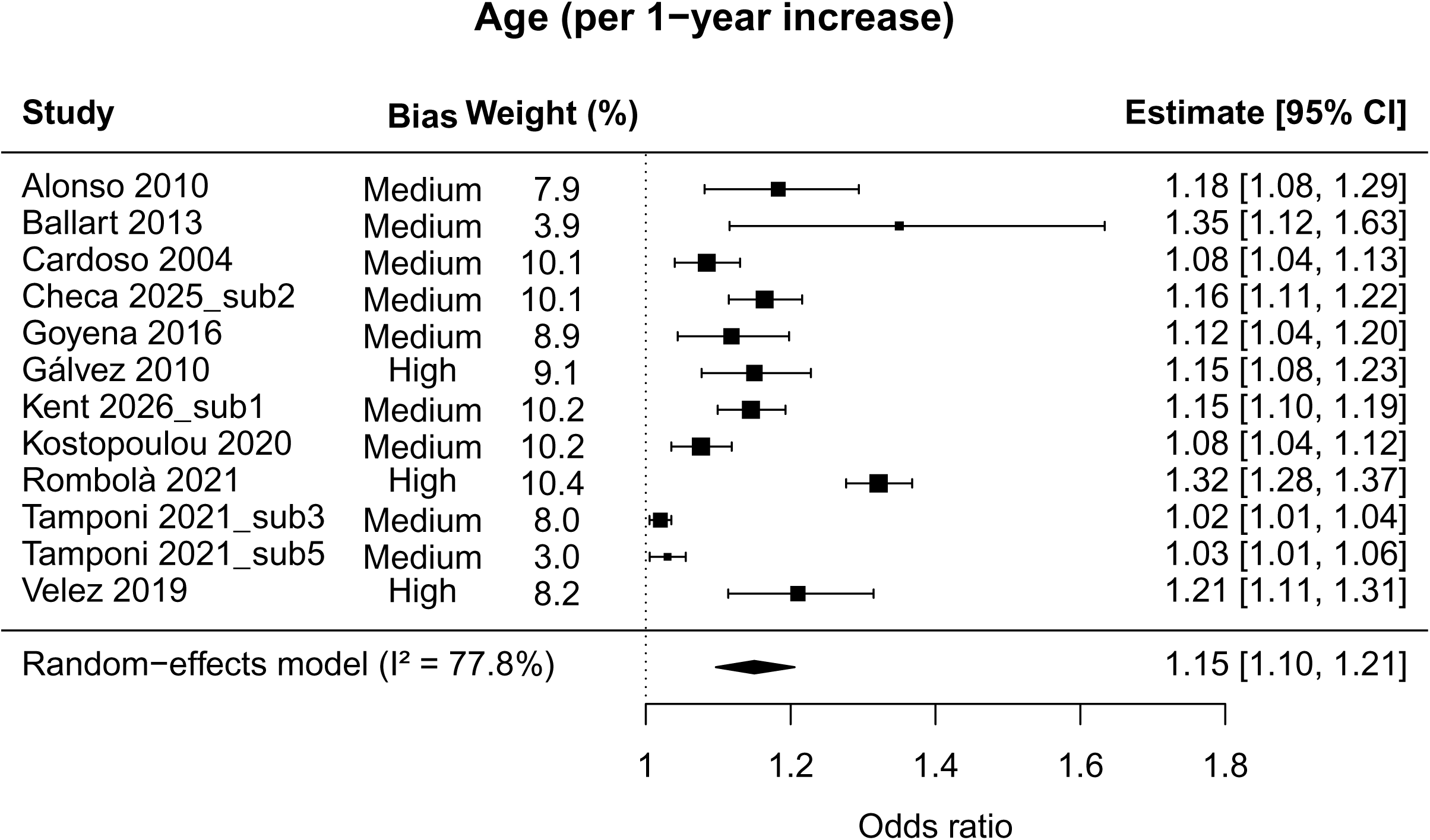
Random-effects forest plot of age (per year) as a risk factor for canine leishmaniosis, based on sub-study specific adjusted odds ratios (OR). Squares represent OR with 95% confidence intervals (CI), with square size proportional to sub-study weight. The diamond represents the pooled summary estimate, while total heterogeneity is quantified by I^2^. The risk of bias for each sub-study is shown.

The secondary analysis, using only unadjusted estimates (n = 30 sub-studies), showed a weaker but still significant association between age and *Leishmania* infection, with a one year increase in age resulting in a 6% increase in the odds of infection (p = 0.002, Table 3, Additional file 2: Fig. S3). There was considerable heterogeneity (I^2^ = 69.7%, Q = 847.8, df = 29, p < 0.001), with all heterogeneity attributable to within-study variance (i.e. between sub-studies nested in studies).

Sensitivity analyses showed mostly similar patterns as the primary and secondary analyses, with older dogs having higher odds of infection than younger dogs (Additional file 2: Appendix, Fig. S4, S5, S6, S7).

**Fig. 4.**
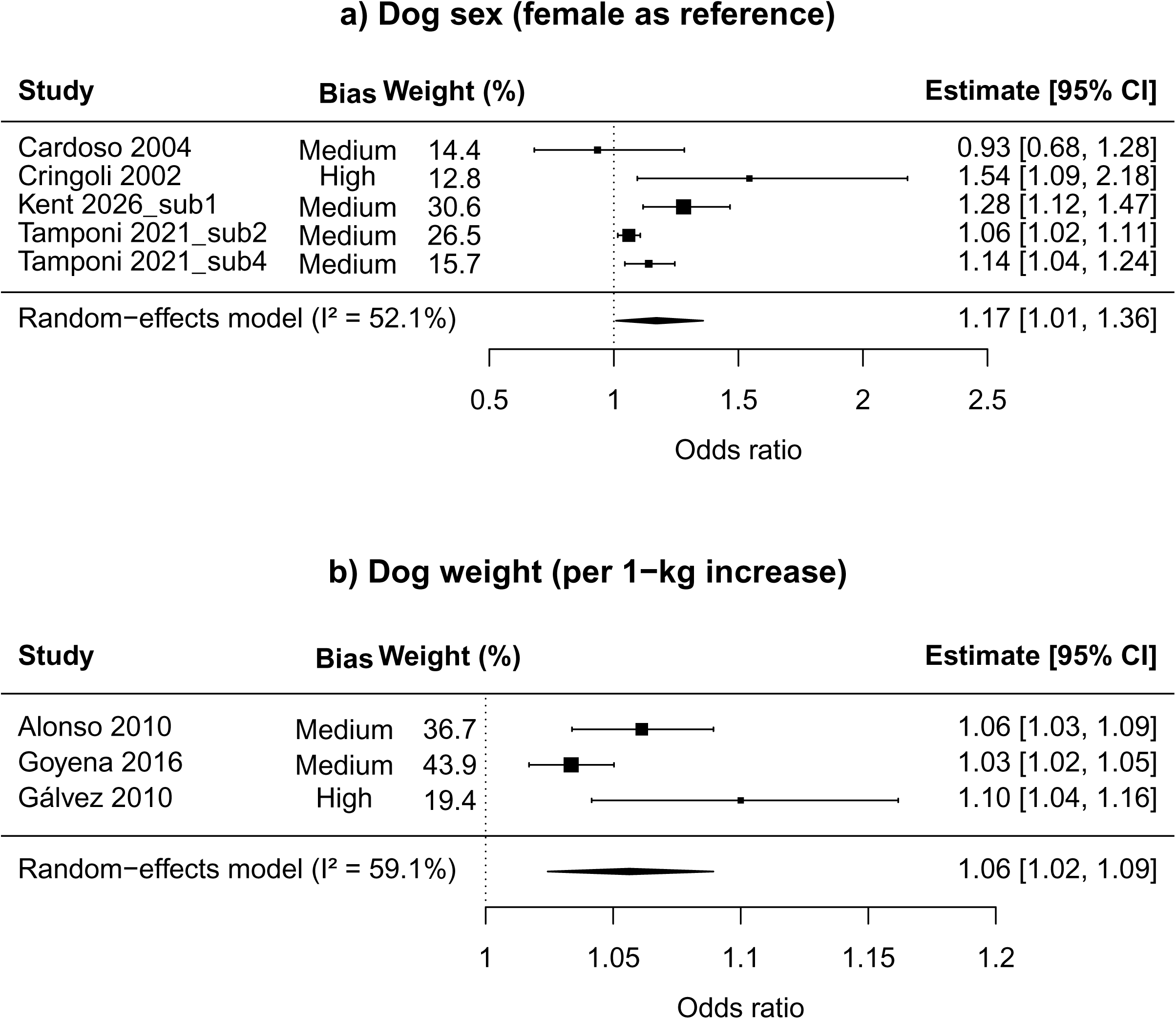
Random-effects forest plots of (**a**) dog sex and (**b**) dog weight as risk factors for *Leishmania* infection in dogs, based on sub-study specific adjusted odds ratios (OR). Squares represent OR with 95% confidence intervals (CI), with square size proportional to sub-study weight. The diamond represents the pooled summary estimate, while total heterogeneity is quantified by I^2^. The dotted line represents OR = 1. The risk of bias for each sub-study is shown.

**Fig. 5.**
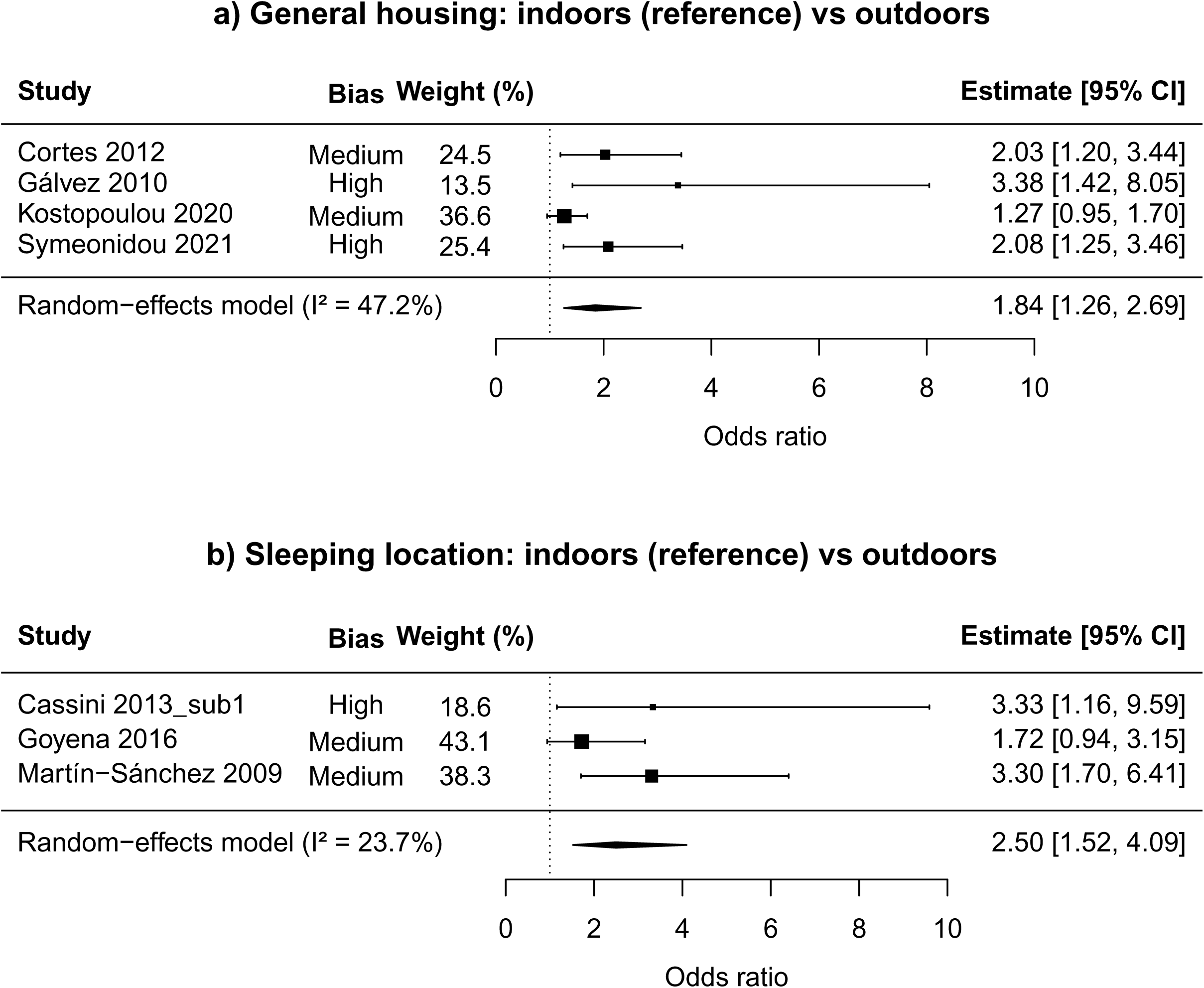
Random-effects forest plots of (**a**) general housing and (**b**) sleeping location as risk factors for *Leishmania* infection in dogs, based on sub-study specific adjusted odds ratios (OR). Squares represent OR with 95% confidence intervals (CI), with square size proportional to sub-study weight. The diamond represents the pooled summary estimate, while total heterogeneity is quantified by I^2^. The dotted line represents OR = 1. The risk of bias for each sub-study is shown.

Overall, there is considerable evidence suggesting that older dogs are at higher risk of infection than younger dogs, but there was also some heterogeneity across studies. This pattern was found across both the primary and secondary analyses, and was largely consistent across sensitivity analyses, even though not all pooled estimates in the sensitivity analyses were statistically significant. Because some studies did not report adjusted age effect sizes due to non significant findings, the pooled estimate from the primary analysis should be interpreted with some caution.

#### Sex of dog

A total of 31 studies (40 sub-studies) with estimates of the association between sex and *Leishmania* infection were included in our final analytic dataset. A significant association was found in the primary analysis, restricted to adjusted estimates (n = 5 sub-studies), with the odds of infection being 17% higher for males compared to females (p = 0.038, Table 2, Fig. 4a). There was evidence of heterogeneity (I^2^ = 52.1%, Q = 12.8, df = 4, p = 0.012), decomposed into 45.6% attributable to between-study variance and 6.5% to between-sub-study (within-study) variance.

A significant association between sex and *Leishmania* infection was also found in the secondary analysis, using only unadjusted estimates (n = 39 sub-studies), with males having 20% higher odds of infection than females (p < 0.001, Table 3, Additional file 2: Fig. S8).

Low heterogeneity was found (I^2^ = 11.7%, Q = 69.0, df = 38, p = 0.002), with all heterogeneity attributable to between-sub-study (within study) variance.

Most sensitivity analyses showed similar patterns as the primary and secondary analyses, with males being at higher odds of infection than females (Additional file 2: Appendix, Fig. S9).

Overall, there is some evidence of males being at higher risk of infection than females. The primary analysis, while consisting of relatively few studies and moderate heterogeneity, showed a significant association. The secondary analysis also found a significant association, and so did most of the sensitivity analyses. However, several studies did not report adjusted effect sizes of sex due to lack of statistical significance, which may reflect reporting bias and reduces confidence in the pooled estimate of the primary analysis.

#### Weight of dog

The final analytic dataset contained estimates of the association between body weight and *Leishmania* infection from a total of 11 studies (12 sub-studies). The primary analysis, restricted to adjusted estimates (n = 3 sub-studies), showed a significant association, with the odds of infection increasing by 6% per one kg increase in weight (p < 0.001, Table 2, Fig. 4b). There was considerable heterogeneity (I^2^ = 59.1%, Q = 6.5, df = 2, p = 0.038), with all heterogeneity attributable to between-study variance.

The secondary analysis, using only unadjusted estimates (n = 9 sub-studies), showed a weaker but still significant association between weight and *Leishmania* infection, with a one kg increase in weight resulting in a 2% increase in the odds of infection (p = 0.003, Table 3, Additional file 2: Fig. S10). There was evidence of heterogeneity (I^2^ = 67.0%, Q = 58.4, df = 8, p < 0.001), with all heterogeneity attributable to between-study variance.

Sensitivity analyses showed somewhat similar patterns as the primary and secondary analyses, with the odds of infection increasing with body weight (Additional file 2: Appendix, Fig. S11, S12, S13, S14, S15).

Overall, there is some evidence of heavier dogs being at higher risk of infection than lighter dogs. Both primary and secondary analyses found a significant association, but both consisted of relatively few studies and high heterogeneity. The sensitivity analyses showed the same trend of increasing risk with increasing weight, but several of them did not reach statistical significance. Because some studies did not report adjusted effect sizes of weight due to lack of statistical significance, the pooled estimate from the primary analysis should be interpreted with caution.

#### General housing

A total of 16 studies (16 sub-studies) with estimates of the association between general housing and *Leishmania* infection were included in our final analytic dataset. A significant association was found in the primary analysis, restricted to adjusted estimates (n = 4 studies), with the odds of infection being 84% higher for dogs living outdoors compared to dogs living indoors (p = 0.002, Table 2, Fig. 5a). There was evidence of heterogeneity (I^2^ = 47.2%, Q = 7.1, df = 3, p = 0.068) between studies.

A significant association between general housing and *Leishmania* infection was also found in the secondary analysis, using only unadjusted estimates (n = 9 sub-studies), with dogs living outdoors having 116% higher odds of infection dogs living indoors (p = 0.010, Table 3, Additional file 2: Fig. S16). Considerable between-study heterogeneity was found (I^2^ = 57.0%, Q = 123.2, df = 8, p < 0.001).

Sensitivity analyses showed similar patterns as the primary and secondary analyses, with the odds of infection being higher in dogs living outdoors compared to dogs living indoors (Additional file 2: Appendix, Fig. S17, S18, S19).

Overall, there is some evidence of dogs living outdoors being at higher risk of infection than dogs living indoors. Both primary and secondary analyses found a significant association, but both consisted of relatively few studies and considerable heterogeneity. The sensitivity analyses showed the same trend of increased risk if living outdoors, but one of them did not reach statistical significance. Because some studies did not report adjusted effect sizes of general housing due to lack of statistical significance, the pooled estimate from the primary analysis should be interpreted with caution.

#### Sleeping location

The final analytic dataset contained estimates of the association between sleeping location and *Leishmania* infection from a total of 7 studies (7 sub-studies). The primary analysis, restricted to adjusted estimates (n = 3 sub-studies), showed a significant association, with the odds of infection being 150% higher for dogs sleeping outdoors compared to dogs sleeping indoors (p < 0.001, Table 2, Fig. 5b). There was low between-study heterogeneity (I^2^ = 23.7%, Q = 2.4, df = 2, p = 0.299).

The secondary analysis, using only unadjusted estimates (n = 7 sub-studies), showed a similar association between sleeping location and *Leishmania* infection, with dogs sleeping outdoors having 116% higher odds of infection than dogs sleeping indoors (p < 0.001, Table 3, Additional file 2: Fig. S20). There was evidence of some heterogeneity (I^2^ = 30.2%, Q = 37.0, df = 6, p < 0.001) between studies.

Sensitivity analyses showed similar patterns as the primary and secondary analyses, with higher odds of infection in dogs sleeping outdoors compared to dogs sleeping indoors (Additional file 2: Appendix, Fig. S21).

Overall, there is some evidence of dogs sleeping outdoors being at higher risk of infection than dogs sleeping indoors. Both primary and secondary analyses consisted of relatively few studies and considerable heterogeneity, but found a significant association. The sensitivity analyses showed the same pattern of increased risk if sleeping outdoors. However, given that adjusted effect sizes for sleeping location were not reported in some studies due to non significant findings, caution is warranted when interpreting the pooled estimate from the primary analysis.

#### Fur length

A total of 11 studies (19 sub-studies) with estimates of the association between fur length and *Leishmania* infection were included in our final analytic dataset. The primary analysis, restricted to adjusted estimates (n = 3 sub-studies), showed that dogs with long hair had 40% lower odds of infection than dogs with short hair, but this association was not statistically significant (p = 0.198, Table 2, Additional file 2: Fig. S22). There was evidence of high heterogeneity (I^2^ = 94.5%, Q = 16.8, df = 2, p < 0.001), decomposed into 94.1% attributable to between-study variance and 0.4% to between-sub-study (within-study) variance.

A significant association between fur length and *Leishmania* infection was found in the secondary analysis, using only unadjusted estimates (n = 14 sub-studies), with dogs in the medium/long fur length category having 16% lower odds of infection than short-haired dogs (p = 0.029, Table 3, Additional file 2: Fig. S23). Low heterogeneity was found (I^2^ = 9.8%, Q = 13.1, df = 13, p = 0.443), with all heterogeneity attributable to between-sub-study (within study) variance.

Sensitivity analyses generally suggest that the odds of infection were lower in longer-haired dogs than in shorter-haired dogs, but not all estimates reached statistical significance (Additional file 2: Appendix, Fig. S24, S25).

Overall, there is weak evidence of longer-haired dogs being at lower risk of infection than shorter-hair dogs. Most analyses, including sensitivity analyses, suggest such an effect, but several of the estimates did not reach statistical significance. The relatively low number of studies, the high heterogeneity in the primary analysis, and the potential for reporting bias (from studies not reporting adjusted estimates due to non-significant findings), all suggest that caution is warranted when interpreting the pooled estimates.

#### Neuter status

The final analytic dataset contained estimates of the association between neuter status and *Leishmania* infection from a total of 3 studies (9 sub-studies). The primary analysis, restricted to adjusted estimates (n = 6 sub-studies), showed a significant association, with the odds of infection being 122% higher for dogs that were neutered than dogs that were entire (p = 0.001, Table 2, Additional file 2: Fig. S26). There was high heterogeneity (I^2^ = 91.2%, Q = 55.7, df = 5, p < 0.001), with all heterogeneity attributable to between-sub-study (within study) variance. Most of this heterogeneity comes from one study[53], where the authors reported a significant interaction between neuter status and dog age, in which the effect of neutering was larger for younger dogs than for older dogs.

The secondary analysis, using only unadjusted estimates (n = 3 studies), showed a similarly significant association between neuter status and *Leishmania* infection, with the odds of infection being 162% higher for dogs that were neutered than dogs that were not (p < 0.001, Table 3, Additional file 2: Fig. S27). There was low between-study heterogeneity (I^2^ = 13.9%, Q = 3.1, df = 2, p = 0.211).

Sensitivity analyses showed similar patterns as the primary and secondary analyses, with the odds of infection being higher in neutered dogs than in entire dogs (Additional file 2: Appendix, Fig. S28).

Overall, there is some evidence of neutered dogs being at higher risk of infection than entire dogs. Both primary and secondary analyses found a significant association, but both consisted of relatively few studies, and the primary analysis had high heterogeneity. The sensitivity analyses showed the same trend of increased risk in neutered dogs. Lastly, given that one study did not report the adjusted effect size for neuter status because of non significant findings, some caution is warranted when interpreting the pooled estimate from the primary analysis.

#### Breed

A total of 19 studies (28 sub-studies) with estimates of the association between dog breed and *Leishmania* infection were included in our final analytic dataset. However, no primary analysis was conducted since fewer than three harmonizable adjusted estimates were available.

Two secondary analyses (using only unadjusted estimates) were conducted. The first analysis, based on 25 sub-studies, found no significant difference in the odds of infection between pure breeds and dogs classified as mixed/mongrel (p = 0.728, Table 3, Additional file 2: Fig. S29). However, considerable heterogeneity was found (I^2^ = 76.5%, Q = 593.4, df = 24, p < 0.001), decomposed into 35.7% attributable to between-study variance and 40.7% to between-sub-study (within-study) variance. On the other hand, the second analysis, based on 12 sub-studies, found a significant association with dogs classified as mixed/pure having 35% higher odds of infection than mongrel dogs (p = 0.011, Table 3, Additional file 2: Fig. S30). Low heterogeneity was observed (I^2^ = 14.4%, Q = 12.4, df = 11, p = 0.333), all attributable to between-study variance.

Sensitivity analyses showed inconsistent results when compared to secondary analyses but partly suggested that pure breeds may have higher odds of infection than other breeds (Additional file 2: Appendix, Fig. S31, S32, S33, S34).

Overall, there is weak and inconsistent evidence of dog breeds differing in their risk of infection. Across analyses, findings were heterogenous and sensitive to modeling assumptions: only one of two secondary analyses found a significant association, indicating lower risk in mongrel dogs. Sensitivity analyses yielded conflicting results regarding the risk of infection, particularly for pure breeds. Some non-harmonized studies reported significant differences between specific breeds, but several did not report adjusted effect sizes of breed when results were non-significant. This inconsistency likely reflects considerable variation in breed composition between studies, where the broad “purebred” and “mixed” categories may capture different breed-specific risk profiles depending on which breeds are included in each study. Taken together, the evidence suggests that breed may be relevant to infection, but the direction and magnitude of the associations remain uncertain and context dependent.

#### Utilization

The final analytic dataset contained estimates of the association between dog utilization and *Leishmania* infection from a total of 11 studies (11 sub-studies). No primary analysis was conducted since fewer than three harmonizable adjusted estimates were available.

Two secondary analyses (using only unadjusted estimates) were conducted. The first analysis, based on 9 sub-studies, found a significant association where non-hunting dogs had 32% lower odds of infection than hunting dogs (p = 0.047, Table 3, Additional file 2: Fig. S35). There was also considerable between-study heterogeneity (I^2^ = 73.3%, Q = 80.6, df = 8, p < 0.001). Moreover, the second analysis, based on 10 sub-studies, found a significant association where non-pet dogs had 50% higher odds of infection than dogs classified as pets (p = 0.005, Table 3, Additional file 2: Fig. S36). Some between-study heterogeneity was observed (I^2^ = 39.1%, Q = 27.3, df = 9, p = 0.001).

Sensitivity analyses were largely consistent with the secondary analyses, showing higher odds of infection for hunting dogs, and lower odds of infection for pets (Additional file 2: Appendix).

Overall, there is weak evidence of dog utilization modifying the risk of infection, with hunting dogs being at higher risk and pets possibly being at lower risk. Both secondary analyses and sensitivity analyses were largely consistent in this assessment, but none were adjusted for potential confounder or modifier variables. The lack of adjusted estimates, and the potential for reporting bias (from studies not reporting adjusted estimates with non-significant findings), suggests that caution is warranted when interpreting the pooled estimates.

#### Urbanicity

A total of 12 studies (13 sub-studies) with estimates of the association between urbanicity and *Leishmania* infection were included in our final analytic dataset. However, no primary analysis was conducted since fewer than three harmonizable adjusted estimates were available.

The secondary analysis, using only unadjusted estimates (n = 6 studies), showed a significant association between urbanicity and *Leishmania* infection, with the odds of infection being 30% lower for dogs living in periurban/urban environments compared to dogs living in rural environments (p = 0.017, Table 3, Additional file 2: Fig. S37). There was low between-study heterogeneity (I^2^ = 16.4%, Q = 12.3, df = 5, p = 0.030).

Sensitivity analyses showed inconsistent results when compared to the secondary analysis, with several analyses showing no significant difference between levels of urbanicity (Additional file 2: Appendix, Fig. S38, S39).

Overall, there is no clear evidence of urbanicity modifying the risk of infection, but a small tendency where higher risk is observed in more rural locations than in urban ones. The secondary analysis and sensitivity analyses were partly in conflict, and neither were adjusted for potential confounder or modifier variables. These limitations, combined with potential reporting bias from studies that excluded urbanicity from adjusted analyses based on non-significant results, mean that the effect of urbanicity remains unclear.

#### Care setting

The final analytic dataset contained estimates of the association between care setting and *Leishmania* infection from a total of 5 studies (6 sub-studies). No primary analysis was conducted since fewer than three harmonizable adjusted estimates were available.

The secondary analysis, using only unadjusted estimates (n = 5 sub-studies), showed a significant association between care setting and *Leishmania* infection, with the odds of infection being 23% lower for stray dogs compared to dogs living at home (p = 0.026, Table 3, Additional file 2: Fig. S40). No heterogeneity was detected (I^2^ = 0.0%, Q = 3.4, df = 4, p = 0.487).

Sensitivity analyses were mostly consistent with the secondary analysis, suggesting that dogs living at home have higher odds of infection than dogs under other care settings (Additional file 2: Appendix, Fig. S41).

Overall, there is weak evidence of dogs living at home being at higher risk of infection than dogs under other care settings. The secondary analysis and sensitivity analyses were largely consistent in this assessment, and so were two non-harmonized adjusted estimates. However, considering the low number of studies and the overall high potential for bias, caution is advised when interpreting the pooled estimates.

#### Living with other dogs

A total of 5 studies (5 sub-studies) with estimates of the association between cohabitation with other dogs and *Leishmania* infection were included in our final analytic dataset.

However, no primary analysis was conducted since fewer than three harmonizable adjusted estimates were available.

A significant association between cohabitation with other dogs and *Leishmania* infection was found in the secondary analysis, using only unadjusted estimates (n = 5 sub-studies), with dogs that did not cohabit with other dogs having 21% lower odds of infection than dogs that did live with other dogs (p = 0.001, Table 3, Additional file 2: Fig. S42). No heterogeneity was detected (I^2^ = 0.0%, Q = 2.0, df = 4, p = 0.744).

Sensitivity analyses were partly consistent with the secondary analysis but were less comprehensive due to low numbers of available studies (Additional file 2: Appendix).

Overall, there is weak evidence that dogs living with other dogs are at higher risk of infection than dogs that do not cohabit with other dogs. The secondary analysis supports this association, and sensitivity analyses were partly consistent. However, given the small number of studies, the overall high potential for bias from only having unadjusted estimates, and the potential of reporting bias (arising from non-reporting of non-significant estimates), the pooled estimates should be interpreted with caution.

#### Clinical signs

The final analytic dataset contained estimates of the association between clinical signs and *Leishmania* infection from a total of 9 studies (15 sub-studies). No primary analysis was conducted since fewer than three harmonizable adjusted estimates were available.

A significant association between the presence of clinical signs and *Leishmania* infection was found in the secondary analysis, using only unadjusted estimates (n = 15 sub-studies), where dogs that did not show clinical signs had 73% lower odds of infection than dogs that did show clinical signs (p < 0.001, Table 3, Additional file 2: Fig. S43).

Considerable heterogeneity was detected (I^2^ = 71.0%, Q = 93.2, df = 14, p < 0.001), decomposed into 43.1% attributable to between-study variance and 27.9% to between-sub-study (within-study) variance.

Sensitivity analyses were mostly consistent with the secondary analysis, suggesting higher odds of infection in dogs with clinical signs (Additional file 2: Appendix).

Overall, there is weak evidence that *Leishmania* infection is more frequent among dogs showing clinical signs than among dogs without clinical signs. This is supported by the secondary analysis and is mostly consistent across sensitivity analyses. However, since no adjusted estimates were available, some caution is advised when interpreting the pooled estimate.

#### Other risk factors

In addition to the risk factors included in the formal meta-analyses, several studies examined other potential risk factors for *Leishmania* infection that could not be harmonized across studies due to limited numbers of comparable estimates. These are summarized below.

##### Density of sand flies

One study[58] reported significantly higher seroprevalence in dogs living in areas with higher sand fly density than in areas with lower sand fly density.

##### Land cover and home features

One study[30] found significantly higher seroprevalence if the ground type was conglomerate dust, there was less water cover, and there was more bushes/woodland than grassland. Another study[66] reported significantly higher seroprevalence if the land cover consisted of more forest. Lasty, a study[51] showed that seroprevalence depended on the home type (e.g. farmland or apartment) and the predominant type of land cover.

##### Co-infection

Six studies reported associations between *Leishmania* infection and co-infections with other pathogens[33, 37, 45, 46, 56, 68]. These showed mixed results, but some studies reported significantly higher *Leishmania* seroprevalence among dogs positive for *Toxoplasma gondii* or *Neospora caninum*.

##### Owner perception/knowledge

One study[35] reported significantly higher seroprevalence among dogs whose owner perceived the dog to be at higher risk of infection. Another study[48] found no association between seroprevalence and dog owners’ knowledge of leishmaniosis.

##### Travel history and origin of dog

Three studies found no significant association between different aspects of travel history (travel to endemic areas[39], national and international travel[46], travel away from home[58]) and seroprevalence. Additionally, one study[73] reported no significant association between dog origin (originating in Germany or not) and seroprevalence.

##### Coat color

One study[57] examined coat color as a potential risk factor across 7 sub-studies. Of these, 6 sub-studies showed no significant association between coat color and *Leishmania* infection, whereas one sub-study reported significantly higher seroprevalence in dogs with brighter coats.

##### Socioeconomic deprivation

One study[53] used the Index of Multiple Deprivation and reported a significantly lower *Leishmania* seroprevalence in the most deprived group.

##### Cohabitation with animals

Two studies[35, 51] found no statistically significant association between *Leishmania* seroprevalence and cohabitation with animals in the broad sense. Similarly, one study[51] reported no association between seroprevalence and living with cats, and another study[30] showed no association between seroprevalence and cohabitation with seropositive dogs.

##### Human-related factors

One study[51] found no statistically significant association between *Leishmania* seroprevalence and the number of people in the household. Another study[62] reported no association between seroprevalence and whether the dog lived in a human leishmaniasis outbreak area.

##### Preventive measures

Factors that relate to the use of repellents, vaccination, owner knowledge of preventives, or similar, were considered out of scope for this review and were not analyzed. Studies that exclusively reported preventive measures were excluded during screening, but 15 studies that reported both preventive and non-preventive risk factors were retained for analysis of the latter.

Overall, evidence for these additional risk factors was limited to a small number of largely non-comparable estimates, preventing pooled analysis and limiting interpretation. No clear conclusion about the role of these factors can be drawn from the available evidence.

### Publication bias

Publication bias was assessed for the five risk factors that had estimates from ≥10 studies. Funnel plots are shown on Additional file 2: Fig. S44. The funnel plot of adjusted estimates of age (per year) suggested some asymmetry, with a significant Egger’s test (p = 0.034) but a non-significant rank correlation test (p = 0.197). No clear sign of asymmetry was observed in the funnel plot of unadjusted age (per year) estimates, and the tests were non-significant (Egger’s test p = 0.728, rank-test p = 0.596). Unadjusted estimates of sex suggested no asymmetry on the funnel plots, and tests were non-significant (Egger’s test p = 0.704, rank-test p = 0.701). No sign of asymmetry was observed in the funnel plot of unadjusted breed (pure vs mixed/mongrel) estimates, and the tests were non-significant (Egger’s test p = 0.084, rank-test p = 0.799). Lastly, unadjusted estimates of utilization (pets vs other) suggested no asymmetry on the funnel plots, and tests were non-significant (Egger’s test p = 0.304, rank-test p = 0.728). Overall, there was no clear evidence of publication bias, except for the adjusted age estimates where the evidence was inconsistent and may reflect high heterogeneity.

## Discussion

This systematic review and meta-analysis provides a comprehensive quantitative synthesis of risk factor associations with *Leishmania* infection in dogs in Europe, contributing to improved understanding of infection determinants, assessing the strength of existing evidence, and identifying key gaps for future research. Of the identified risk factors, dog age was the only risk factor with considerable evidence of an association with infection, where older dogs were found to have a higher risk of infection than younger dogs. Next followed five risk factors with some evidence, namely sex (males at higher risk), weight (heavier dogs at higher risk), general housing and sleeping location (dogs living/sleeping outdoors at higher risk), and neuter status (neutered dogs at higher risk). Additionally, there was weak evidence of six risk factors (fur length, breed, utilization, care setting, cohabiting with other dogs, and clinical signs) being associated with infection. Other risk factors showed no clear evidence of an association, either due to inconsistent findings or because of few studies quantifying the risk. Compared to reviews conducted in other regions[9–12, 15], both similarities and differences were observed. Exposure-related factors, particularly housing conditions such as outdoor sleeping, showed associations in the Americas that are consistent with our estimates. Age was also consistently identified as a risk factor across regions, but the strength of evidence varied. In contrast, findings of sex as a risk factor showed either weaker or absent associations in other regions compared to in our review. Notably, neuter status emerged as a potential risk factor in our analysis but has not been considered in other regions. While some additional risk factors showed partial consistency across reviews, others were either not assessed or lacked sufficient evidence for comparison. Overall, these regional differences likely reflect differences in ecological conditions, cultural practices (e.g. neutering practices), study methodology, and availability of evidence.

Host-related factors (age, sex, weight and breed) had variable levels of support as risk factors, ranging from weak to considerable evidence. Increasing age was consistently associated with higher risk of *Leishmania* infection. While this may reflect age-related immune changes, it is more likely explained by cumulative exposure, since most studies measured seroprevalence[75]. Older dogs have been exposed to sand fly bites for longer time and may remain chronically infected, leading to an accumulation of seropositive dogs with age. Being a male dog was associated with higher risk of infection, which authors have argued may relate to testosterone reducing immunity to the *Leishmania* parasite, or due to males roaming more and therefore being more exposed to sand flies[53]. Additionally, higher body weight was associated with a higher risk. This may partly relate to heavier dogs emitting more CO_2_ and other bodily odors, which act as olfactory cues for sand flies[76].

However, it is more likely to reflect that heavier dogs, which are often proxies for heavier breeds, are more commonly used for outdoors activities such as guarding and hunting, thereby increasing their exposure to sand flies[75]. As to breeds, the inconsistent findings indicate that any association is context dependent. Some breeds may be more genetically susceptible to infection (e.g. Boxers and Rottweilers[77]) or be more exposed to sand flies via their typical utilization (e.g. if used for hunting). Broad breed classifications may therefore obscure more specific breed-level effects.

Regarding clinical signs and co-infections, results ranged from weak to no clear evidence. For clinical signs, the observed association with *Leishmania* infection likely reflects reverse causation rather than clinical signs being a risk factor for infection. Clinical signs may be a consequence of infection, such that infected dogs are more likely to develop signs compatible with canine leishmaniosis. Alternatively, clinical signs could represent underlying vulnerability or comorbidity, where dogs with poor health are more likely to also become infected by *Leishmania*. Moreover, clinical signs could be a downstream consequence of *Leishmania* infection, where infected dogs are more likely to develop secondary conditions. Lastly, while there was no clear evidence of an association between *Leishmania* infection and co-infections, such concurrent infections are biologically plausible in dogs with weakened immune systems.

Most of the risk factors identified in this review relate primarily to exposure, by increasing or decreasing the probability of being exposed to sand fly bites. The evidence for these risk factors ranged from some to no clear evidence. Risk factors that directly relate to time spent outdoors (general housing and sleeping location) showed the strongest evidence of an association with *Leishmania* infection. Next followed some risk factors that indirectly relate to exposure, such as fur length (higher likelihood of being bitten if short fur), utilization (e.g. hunting dogs spending more time outdoors than pets), and cohabitation with other dogs (multiple hosts likely attract more sand flies[78]). Lastly, several risk factors showed no clear evidence of an association, either due to inconsistent findings or because of too few studies to draw conclusions. For urbanicity, rural habitats may be more suitable for sand flies[7] and dogs may spend more time outdoors than in urban regions[79], yet findings remained inconsistent. Factors such as sand fly density, land cover and home features likely relate to habitat preferences and exposure to sand flies but had limited study coverage.

Similarly, differences between dogs with different coat colors likely reflect that brighter colors may attract sand flies[57], but evidence remains sparse. Living with other animals or multiple humans may also increase sand fly attraction, but this has also only rarely been examined. Overall, factors such as general housing or sleeping location appear as primary proxies of exposure, while fur length, utilization and cohabitation with other dogs may represent more indirect secondary proxies of exposure.

Evidence for risk factors related to management (neuter status and care setting) ranged from some to weak evidence of an association with *Leishmania* infection. For neuter status, the strongest evidence of neutered dogs being at higher risk came from two studies from the United Kingdom (UK)[41, 53], and the authors propose that this may partly reflect active neutering policies for dogs that are being rehomed or imported to the UK[53]. If imported or rehomed dogs are more frequently neutered than local dogs, neuter status may act as a proxy of previous residence in endemic regions and thus a higher probability of prior infection. Regarding care setting, there was weak evidence suggesting that dogs at home have higher seroprevalence than stray dogs. Several mechanisms may contribute to this pattern. If there are differences in age structure, with stray dogs being younger than owned dogs, this difference may result in higher seroprevalence in the older population. Additionally, survival bias may play a part, if infected stray dogs are more likely to die before sampling compared to domestic dogs with better access to veterinary care, resulting in an underrepresentation of infected stray dogs. Lastly, detection bias, where stray dogs are tested less often, is likely less important here since within-study comparisons largely control for this.

Only a few studies looked at travel history, dog origin, or the owner’s socioeconomics (deprivation, owner perception and knowledge), making it difficult to draw firm conclusions. Travel history to, and origin from, endemic regions are often suggested as possible risk factors, but none of the studies found a significant association with *Leishmania* infection.

This may reflect limitations in how travel is measured, as it is often measured crudely without accounting for duration, timing, specific destinations or intensity of exposure. Additionally, no potential confounders were addressed in the studies, and reverse causation may occur if infected or clinically affected dogs are less likely to travel. Together, these issues likely dilute any true effect, suggesting that travel history is a challenging variable to use as a proxy for risk and may be difficult to interpret reliably. As to socioeconomic deprivation, one study[53] reported lower seroprevalence in the most deprived group. The authors suggest this may reflect lower likelihood of testing and diagnosis among dogs owned by individuals in more deprived settings. Lastly, studies considering owner perception and knowledge of the disease showed higher seroprevalence when owners perceived their dogs to be at higher risk, but no association with owner knowledge. Owners perceiving higher risk likely live in areas with greater local exposure and may already have experience with the disease, leading to increased awareness. In contrast, knowledge of the disease does not necessarily translate into use of preventive measures, and any true effect may be diluted by a combination of knowledgeable owners applying prevention and owners only acquiring knowledge after infection.

A major limitation was inconsistent and inadequate handling of confounding across studies, possibly resulting in reporting bias. While several studies conducted multivariable analyses, many studies did not determine and justify in advance which variables to adjust for or adequately report adjusted estimates. Several studies employed problematic variable selection procedures such as stepwise variable selection or excluding variables based on univariable p-values, which are known to give inflated effect estimates and lead to invalid confidence intervals and p-values[21, 22]. Even when adjusted, inconsistent covariate selection across studies makes synthesis challenging. For example, some studies included age while others did not, meaning that effect estimates may not directly compare and may be subject to residual confounding. In addition, many studies were descriptive and only reported univariable results, which while important during initial investigations, should be interpreted cautiously. We recommend the following to improve future work: (1) pre-specify a core set of confounders based on causal reasoning and existing literature (this review provides a good starting point for risk factors to consider, e.g. age, sex, and sleeping location); (2) report both unadjusted and adjusted estimates with 95% confidence intervals; (3) avoid automatic variable selection and instead specify models based on subject-matter knowledge; (4) try to avoid post-hoc variable exclusion, and justify such exclusions if required due to e.g. missing data or collinearity; (5) ensure that sample size is adequate for the planned multivariable model, or alternatively restrict adjusted models to a pre-specified set of the most relevant confounders when sample size is limited.

Additional limitations relate to study design, outcome definition, and the review process. Most included studies were cross-sectional and observational, meaning that causal inference is limited. Additionally, the most studied outcome was seroprevalence, which captures both past and current infections and may therefore not reflect recent transmission. This can result in a temporal mismatch between measured risk factors and the timing of infection, with age being a key example. Since infections may have occurred at different times prior to sampling, age at sampling may not reflect the age at which the infection was acquired. With respect to the review process, screening, data extraction and risk of bias assessment were conducted by a single reviewer, which may introduce the risk of selection or extraction bias. Moreover, the search strategy was limited to a single database (PubMed), meaning that some relevant studies may have been missed. However, PubMed provides comprehensive indexing of published veterinary and epidemiological literature, and our exclusion of grey literature was applied consistently. Furthermore, substantial variable harmonization was required to compare across studies, including making assumptions about the linearity between risk and age/weight when harmonizing those to a continuous scale, which may have introduced some misclassification or loss of detail. This was, however, mitigated through sensitivity analyses, where multiple alternative codings of variables were considered.

## Conclusions

This review identifies key risk factors for canine *Leishmania* infection that warrant consideration in future research and disease control efforts. Age emerged as the primary risk factor, while sex, body weight, and outdoor living/sleeping represent secondary but relatively consistent associations. Future studies should routinely adjust for these core risk factors in analyses conducted using models with pre-specified covariates informed by subject-matter knowledge, and avoid variable selection procedures that may bias estimates. Beyond methodological improvements, important gaps remain. The relevance of certain risk factors, particularly neuter status, seems to differ between endemic and non-endemic regions, highlighting the need to account for regional context in future studies. Additionally, to strengthen causal inference, future research should prioritize longitudinal cohort designs to better capture causality. For modifiable risk factors (e.g. sleeping location), experimental approaches such as randomized controlled trials should be considered where it is feasible to provide stronger evidence of causality. These findings have direct implications for disease control and prevention. Strategies targeting dogs with the highest risk profiles, such as older dogs living or sleeping outdoors, could improve the effectiveness of interventions.

## Abbreviations

CI: Confidence interval
I^2^: I-squared statistic (measure of heterogeneity)
IFAT: Indirect fluorescent antibody test
OR: Odds ratio
PCR: Polymerase chain reaction
PRISMA: Preferred reporting items for systematic reviews and meta-analyses
REML: Restricted maximum likelihood
ROBINS-E: Risk of bias in non-randomized studies of exposures

## Declarations

### Ethics approval and consent to participate

Ethical approval was not required for this study, as it is based on previously published data.

### Consent for publication

Not applicable.

### Availability of data and material

The datasets generated and/or analyzed during the current study are available in the Zenodo repository, https://doi.org/10.5281/zenodo.21159338. The datasets are also provided as supplementary materials, with sub-study characteristics in Additional file 3: Dataset S1 and effect sizes in Additional file 4: Dataset S2. The R scripts used for data harmonization and meta-analysis are publicly available on GitHub, https://github.com/erlendfossen/Canine_leish_meta.

### Authors’ contributions

Conceptualization: E.I.F.F., C.M., K.A.; Data curation: E.I.F.F.; Formal analysis: E.I.F.F.; Funding acquisition: K.A., C.M.; Investigation: E.I.F.F.; Methodology: E.I.F.F.; Resources: Not applicable; Software: E.I.F.F.; Supervision: K.A.; Visualization: E.I.F.F.; Writing-original draft: E.I.F.F.; Writing-review & editing: All authors. All authors approved the submission of this manuscript.

## Acknowledgements

Not applicable.

