## Additional file 2 for "A systematic review and meta-analysis of risk factors for *Leishmania* infection in dogs in Europe"

### Appendix: sensitivity analyses

#### 1. Age

Sensitivity analyses showed mostly similar patterns as the primary and secondary analyses, with older dogs being at higher odds of infection than younger dogs. Including the risk of bias as a covariate in the primary analysis showed a significantly higher (p = 0.022) odds of infection in studies where the bias was high (pooled OR = 1.23, 95% CI: 1.15-1.33, p < 0.001), compared to studies with medium risk of bias (pooled OR = 1.12, 95% CI: 1.07-1.17, p < 0.001). Fixed-effects models of both the primary and secondary analyses showed results with similar direction, magnitude and significance (**Table 2**, **Table 3**). The mixed-estimates analysis (n = 13 studies / 15 sub-studies) likewise showed a significant association between age (per year) and infection (pooled OR = 1.13, 95% CI: 1.08-1.18, p < 0.001, **Figure S4**), and high total heterogeneity (I^2^ = 75.5%, p <0.001, all attributable to between-study variance). Analyses using different age coding, also showed that older dogs in general were at higher risk than younger dogs: dogs <1 year showed lower odds of infection than dogs ≥1 year (p = 0.019, **Table 3**, **Figure S5**); unadjusted analysis (n = 11 studies / 13 sub-studies) showed that dogs aged 3-6 years were at higher odds of infection than dogs <3 years (OR = 1.75, 95% CI: 1.25-2.44, p = 0.001, **Figure S6**); dogs ≤ 7 year showed a lower, but non-significant, odds of infection than dogs >7 year (p = 0.113, **Table 3**, **Figure S7**). Lastly, non-harmonized studies reporting adjusted estimates (n = 4, all statistically significant) or unadjusted estimates (n = 12, 8 statistically significant), all showed the same pattern with older dogs being at higher risk than younger dogs. A total of 7 studies (8 sub-studies) presented multivariable analyses, but excluded age from the analysis because the effect size did not pass the significance level.

#### 2. Sex

Most sensitivity analyses showed similar patterns as the primary and secondary analyses, with males being at higher odds of infection than females. There was no significant difference (p = 0.137) between studies with medium risk of bias (pooled OR = 1.13, 95% CI: 0.98-1.29, p = 0.083) and studies with high risk of bias (pooled OR = 1.54, 95% CI: 1.04-2.29, p = 0.030) in the primary analysis. Fixed-effects models of both the primary and secondary analyses showed slightly lower OR than random-effects models, but were still significant, with males being at higher risk than females (**Table 2**, **Table 3**). The mixed-estimates analysis (n = 12 studies / 14 sub-studies) likewise showed a significant association with males at higher risk (pooled OR = 1.17, 95% CI: 1.05-1.30, p = 0.004, **Figure S9**), and had low heterogeneity (I^2^ = 10.8%, p = 0.099, decomposed into 10.2% attributable to between-study variance and 0.7% to within-study (between sub-study) variance). However, most of the relevant non-harmonized studies showed a non-significant association between sex and infection. Of non-harmonized studies that reported adjusted estimates, 16 studies (17 sub-studies) of 18 studies (19 sub-studies) excluded sex from their multivariable analyses based on effect sizes not passing significance levels, while two studies showed a significant association with males at higher risk. Of the five non-harmonized studies that reported unadjusted estimates, 1 showed a significant association with males at higher risk, and 4 showed no significant association.

#### 3. Weight

Sensitivity analyses showed somewhat similar patterns as the primary and secondary analyses, with the odds of infection increasing with body weight. There was no significant difference (p = 0.055) between studies with medium risk of bias (pooled OR = 1.04, 95% CI: 1.02-1.06, p < 0.001) and studies with high risk of bias (OR = 1.10, 95% CI: 1.04-1.16, p < 0.001) in the primary analysis. Fixed-effects models showed slightly lower OR than random-effects models, but the same direction and significance, with the odds of infection increasing with weight (**Table 2**, **Table 3**). The mixed-estimates analysis (n = 4 studies / 4 sub-studies) showed a significant association between weight (per year) and infection (pooled OR = 1.04, 95% CI: 1.01-1.08, p = 0.017, **Figure S11**), and high total heterogeneity (I^2^ = 76.6%, p = 0.004). Analyses using different weight coding, showed that heavier dogs in general were at higher risk than lighter dogs, but the pooled estimates were not always statistically significant: dogs >25 kg showed higher odds of infection than dogs <25 kg (p = 0.177, **Table 3**, **Figure S12**); dogs categorized as large showed significantly higher odds of infection than dogs categorized as small (p = 0.006, **Table 3**, **Figure S13**); unadjusted analyses (n = 5 studies / 6 sub-studies) showed that dogs categorized as medium had a higher odds of infection than dogs categorized as small (OR = 1.38, 95% CI: 0.93-2.05, p = 0.115, **Figure S14**), and dogs categorized as large had a significantly higher odds than dogs categorized as medium (OR = 1.41, 95% CI: 1.02-1.97, p = 0.039, **Figure S15**). Lastly, of non-harmonized studies that reported adjusted estimates, 5 studies (6 sub-studies) excluded body weight from their multivariable analyses based on effect sizes not passing significance levels. Of the two non-harmonized studies that reported unadjusted estimates, both showed that heavier dogs were at higher risk, but neither estimate reached statistical significance.

#### 4. General housing

Sensitivity analyses showed similar patterns as the primary and secondary analyses, with the odds of infection being higher in dogs living outdoors compared to dogs living indoors. Including the risk of bias as a covariate in the primary analysis showed a significantly higher (p = 0.048) odds of infection in studies where the bias was high (pooled OR = 2.36, 95% CI: 1.52-3.65, p < 0.001), compared to studies with medium risk of bias (pooled OR = 1.42, 95% CI: 1.10-1.83, p = 0.008). Fixed-effects models showed slightly lower OR than random-effects models, but with the same direction and significance (**Table 2**, **Table 3**). Similarly, the mixed-estimates analysis (n = 6 studies / 6 sub-studies) showed a significant association with outdoor dogs being at higher odds of infection than indoor dogs (pooled OR = 1.62, 95% CI: 1.05-2.49, p = 0.028, **Figure S17**), and some heterogeneity (I^2^ = 40.1%, p < 0.001). Analyses where different coding was used for general housing, showed that living more outdoors than indoors is associated with higher odds of infection, but the pooled estimates were not always statistically significant: unadjusted analyses (n = 7 studies / 7 sub-studies) showed that dogs living partly or fully outdoors had significantly higher odds of infection than dogs living exclusively indoors (OR = 1.72, 95% CI: 1.04-2.85, p = 0.035, **Figure S18**), and that dogs living fully outdoors had higher odds than dog living partly outdoors (OR = 1.39, 95% CI: 0.92-2.10, p = 0.119, **Figure S19**). Lastly, 6 non-harmonized studies (6 sub-studies) reported adjusted estimates, but excluded general housing from their multivariable analyses based on effect sizes not passing significance levels.

#### 5. Sleeping location

Sensitivity analyses showed similar patterns as the primary and secondary analyses, with higher odds of infection in dogs sleeping outdoors compared to dogs sleeping indoors. There was no significant difference (p = 0.535) between studies with medium risk of bias (pooled OR = 2.32, 95% CI: 1.48-3.62, p < 0.001) and studies with high risk of bias (OR = 3.33, 95% CI: 1.16-9.59, p < 0.026) in the primary analysis. Fixed-effects models showed slightly lower OR than random-effects models, but the same direction and significance, with the odds of infection being higher in dogs sleeping outdoors (**Table 2**, **Table 3**). Similarly, the mixed-estimates analysis (n = 6 studies / 6 sub-studies) showed a significant association where dogs sleeping outdoors had higher odds of infection than dogs sleeping indoors (pooled OR = 2.00, 95% CI: 1.24-3.22, p = 0.005, **Figure S21**), and some total heterogeneity (I^2^ = 38.8%, p = 0.002). Lastly, of the 5 non-harmonized studies (5 sub-studies) that reported adjusted estimates, 1 showed that dogs sleeping outdoors had higher odds of infection, while the remaining 4 studies excluded sleeping location from their multivariable analyses because effect sizes did not pass significance levels.

#### 6. Fur length

Sensitivity analyses generally suggest that the odds of infection were lower in longer-haired dogs than in shorter-haired dogs, but not all estimates reached statistical significance. All studies in the primary analysis had moderate risk of bias. Fixed-effects models of the primary and secondary analyses had similar estimates (OR ~ 0.85) and were both statistically significant (**Table 2, Table 3**). The mixed-estimates analysis (n = 3 studies / 5 sub-studies) showed a non-significant association with long-haired dogs having lower odds of infection than short-haired dogs (pooled OR = 0.59, 95% CI: 0.34-1.04, p = 0.069, **Figure S24**), and high heterogeneity (I^2^ = 83.2%, p < 0.001, 82.0% attributable to between-study variance and 1.2% to between-sub-study variance). When considering different coding of fur length for the secondary analysis, unadjusted analyses (n = 3 studies / 4 sub-studies) showed a non-significant association with long-haired dogs having similar odds as the reference category, short-haired dogs (OR = 1.04, 95% CI: 0.92-1.17, p = 0.554, **Figure S25**). Lastly, one study reported a non-significant unadjusted estimate, but was not possible to harmonize, whereas 4 non-harmonized studies (5 sub-studies) reported adjusted estimates, but excluded fur length from their multivariable analyses based on effect sizes not passing significance levels.

#### 7. Neuter status

Sensitivity analyses showed similar patterns as the primary and secondary analyses, with the odds of infection being higher in neutered dogs than in entire dogs. All studies in the primary analysis had moderate risk of bias. Fixed-effects models gave very similar pooled estimates as the random-effects models, with the odds of infection being higher in neutered dogs (**Table 2**, **Table 3**). The mixed-estimates analysis (n = 3 studies / 7 sub-studies) also showed a similar and significant association where neutered dogs had higher odds than entire dogs (pooled OR = 2.15, 95% CI: 1.39-3.31, p < 0.001, **Figure S28**), and there was considerable total heterogeneity (I^2^ = 85.6%, p < 0.001, all attributable to between-sub-study variance). Lastly, one non-harmonized study reported adjusted estimates, but excluded reporting of neuter status from their adjusted analysis based on the effect size not reaching statistical significance.

#### 8. Breed

Sensitivity analyses showed inconsistent results when compared to the secondary analyses. All studies in secondary analyses had high risk of bias. Fixed-effects models showed similar OR as random-effects models, but with narrower confidence intervals (**Table 3**). The mixed-estimates analysis (n = 5 studies / 7 sub-studies) showed lower odds of infection in mixed/mongrel breeds than in pure breeds, but the estimate was non-significant (pooled OR = 0.87, 95% CI: 0.67-1.13, p = 0.308, **Figure S31**), and with considerable heterogeneity (I^2^ = 71.2%, p < 0.001). Analyses using different coding of breed categories indicated that pure breeds may have higher odds of infection: unadjusted analysis (n = 3 studies / 9 sub-studies) showed that mixed breeds had lower odds of infection than pure breeds (OR = 0.72, 95% CI: 0.53-0.97, p = 0.032, **Figure S32**); unadjusted analysis (n = 4 studies / 10 sub-studies) showed that mongrel dogs had lower odds of infection than pure breeds (OR = 0.71, 95% CI: 0.53-0.95, p = 0.021, **Figure S33**); unadjusted analysis (n = 3 studies /9 sub-studies) showed that mixed breeds had similar odds of infection as mongrel dogs (reference category) (OR = 1.09, 95% CI: 0.67-1.79, p = 0.787, **Figure S34**). Lastly, non-harmonized studies reporting adjusted estimates (n = 2, both statistically significant) or unadjusted estimates (n = 8, 5 statistically significant), showed that different specific breeds had higher or lower risk of infection than other breeds. A total of 12 studies (13 sub-studies) presented multivariable analyses, but excluded breed from the analysis because the effect size did not reach significance.

#### 9. Utilization

Sensitivity analyses were largely consistent with the secondary analyses, showing higher odds of infection for hunting dogs, and lower odds of infection for pets. All studies in the secondary analysis had high risk of bias. Fixed-effects models gave similar pooled estimates as the random-effects models, with the odds of infection being higher in hunting dogs and lower in pets (**Table 3**). Not enough harmonizable adjusted estimates were available for a mixed-estimates analysis. Lastly, two non-harmonized studies reported statistically significant adjusted estimates where hunting dogs were at higher risk than pets. Additionally, 4 non-harmonized studies reporting adjusted estimates, and one study reporting unadjusted estimates, did not report effect sizes because they were statistically non-significant.

#### 10. Urbanicity

Sensitivity analyses showed inconsistent results when compared to the secondary analysis. All studies in secondary analyses had high risk of bias. The fixed-effects model agreed with the random-effects model (**Table 3**). Not enough harmonizable adjusted estimates were available for a mixed-estimates analysis. Analyses using different coding of urbanicity categories, all using unadjusted estimates, showed no difference between periurban dogs (reference category) and rural dogs (n = 6 studies / 6 sub-studies, OR = 0.94, 95% CI: 0.80-1.11, p = 0.478, **Figure S38**), and no difference between urban (reference) and rural dogs (n = 4 studies / 4 sub-studies, OR = 0.95, 95% CI: 0.47-1.93, p = 0.894, **Figure S39**). Lastly, among the 8 non‑harmonized studies (9 sub‑studies) that reported adjusted estimates, statistically significant associations were observed in 3 studies (4 sub‑studies). Of these, two studies identified rural dogs as being at higher risk, while one study (two sub‑studies) reported higher risk in periurban dogs compared with both urban and rural dogs. The remaining five studies did not include adjusted estimates for urbanicity because results were non‑significant.

#### 11. Care setting

Sensitivity analyses were mostly consistent with the secondary analysis, suggesting that dogs living at home are at higher infection risk than dogs under other care settings. All studies in secondary analyses had high risk of bias. The fixed-effects model yielded the same pooled estimate as the random-effects model, since the detected heterogeneity was zero (**Table 3**). Not enough harmonizable adjusted estimates were available for a mixed-estimates analysis. Comparing dogs living at home to a broader category of care settings, showed no significant difference between dogs at home (reference) and dogs under other care settings (n = 3 studies / 4 sub-studies, OR = 1.02, 95% CI: 0.14-7.48, p = 0.984, **Figure S41**). Lastly, two non-harmonized studies reported statistically significant adjusted estimates, with owned dogs being at higher risk than kennel dogs, and with sheltered dogs being at lower risks than both owned and stray dogs. Additionally, three non-harmonized studies (4 sub-studies) reported statistically significant unadjusted estimates showing that owned dogs were at higher risk than kennel dogs, and that kennel dogs were at higher risk than farm dogs.

#### 12. Living with other dogs

Sensitivity analyses were partly consistent with the secondary analysis. All studies in secondary analysis had high risk of bias. The fixed-effects model yielded the same pooled estimate as the random-effects model, since the detected heterogeneity was zero (**Table 3**). Not enough harmonizable adjusted estimates were available for a mixed-estimates analysis. Lastly, 6 non-harmonized studies reported adjusted estimates, but did not report adjusted estimates of cohabitation with other dogs, due to the effect size not reaching statistical significance.

#### 13. Clinical signs

Sensitivity analyses were mostly consistent with the secondary analysis. All studies in secondary analysis had high risk of bias. The fixed-effects model showed a similar pooled estimate as the random-effects model (**Table 3**). Not enough harmonizable adjusted estimates were available for a mixed-estimates analysis. Lastly, of the 4 non-harmonized studies that reported adjusted estimates, one study showed statistically significantly higher odds of infection if the dog had clinical signs, while 3 studies did not report adjusted estimates because the effect size did not reach statistical significance.

### Supplementary figures

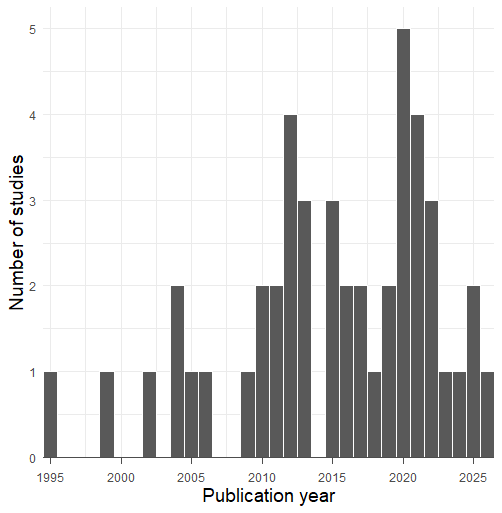

**Figure S1.** Histogram showing the publication year of included studies (n = 46).

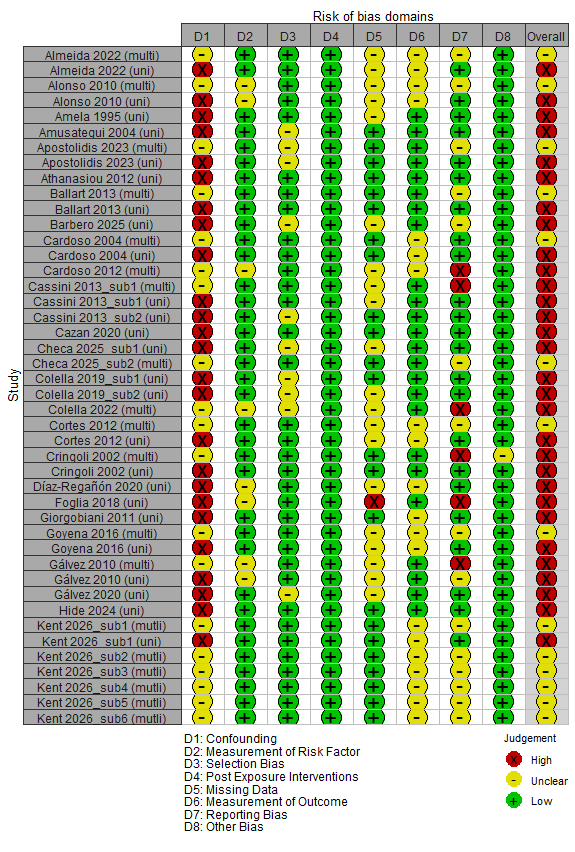

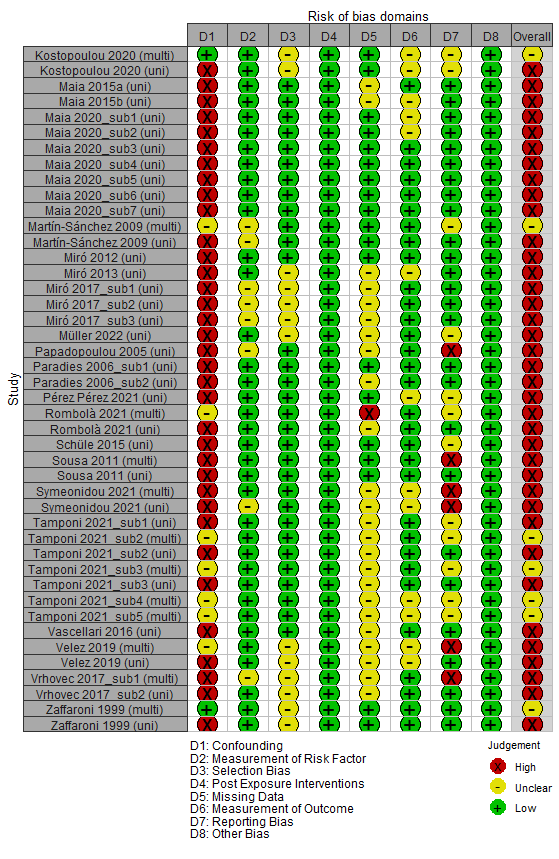

**Figure S2.** Risk of bias across ROBINS-E domains for each analytical unit within sub-studies (figure split across pages for readability). The labels “uni” and “multi” appended to each sub-study ID indicate if the analytical unit was based on reported univariable (unadjusted) or multivariable (adjusted) analyses.

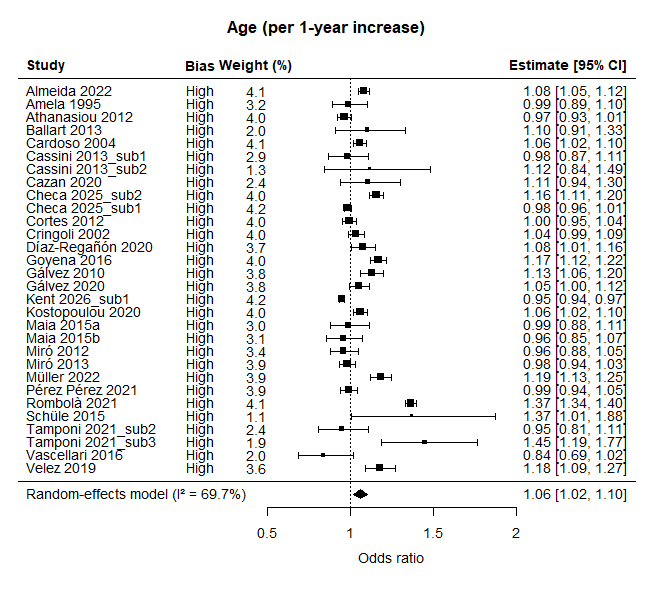

**Figure S3.** Random-effects forest plot of age (per year) as a risk factor for *Leishmania* infection in dogs, based on sub-study specific *unadjusted* odds ratios (OR). All forest plots in Additional file 2 share a common structure: squares represent OR with 95% confidence intervals (CI), with square size proportional to sub-study weight. The diamond represents the pooled summary estimate, while total heterogeneity is quantified by I^2^. The dotted line represents OR = 1. The risk of bias for each sub-study is shown.

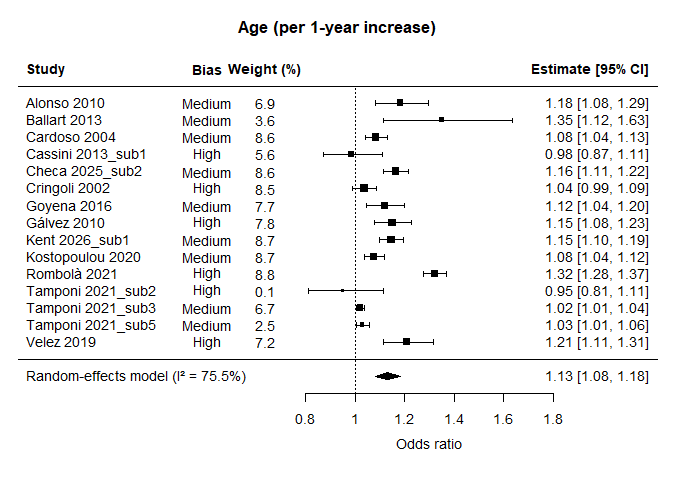

**Figure S4.** Random-effects forest plot of age (per year) as a risk factor for *Leishmania* infection in dogs, based on sub-study specific *mixed-estimates* odds ratios. Plot elements are defined in Figure S3.

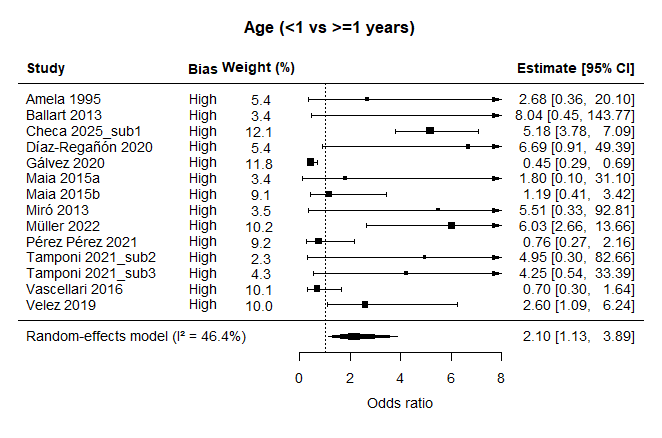

**Figure S5.** Random-effects forest plot of age (<1 vs ≥1 years old) as a risk factor for *Leishmania* infection in dogs, based on sub-study specific *unadjusted* odds ratios. Plot elements are defined in Figure S3.

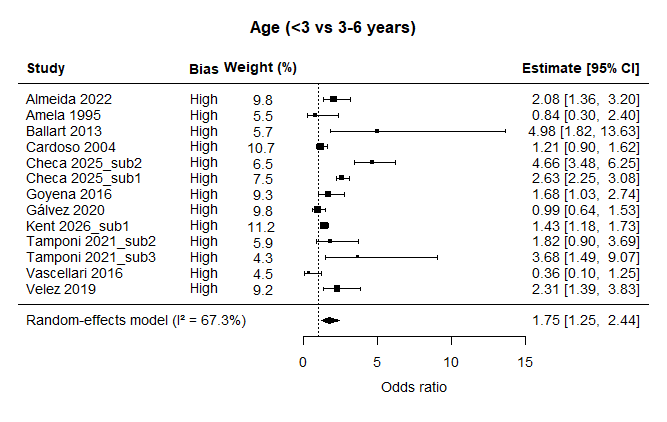

**Figure S6.** Random-effects forest plot of age (<3 vs 3-6 years old) as a risk factor for *Leishmania* infection in dogs, based on sub-study specific *unadjusted* odds ratios. Plot elements are defined in Figure S3.

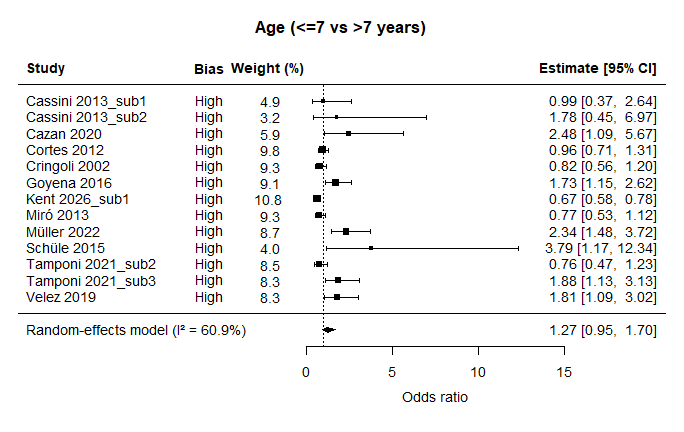

**Figure S7.** Random-effects forest plot of age (≤7 vs >7 years old) as a risk factor for *Leishmania* infection in dogs, based on sub-study specific *unadjusted* odds ratios. Plot elements are defined in Figure S3.

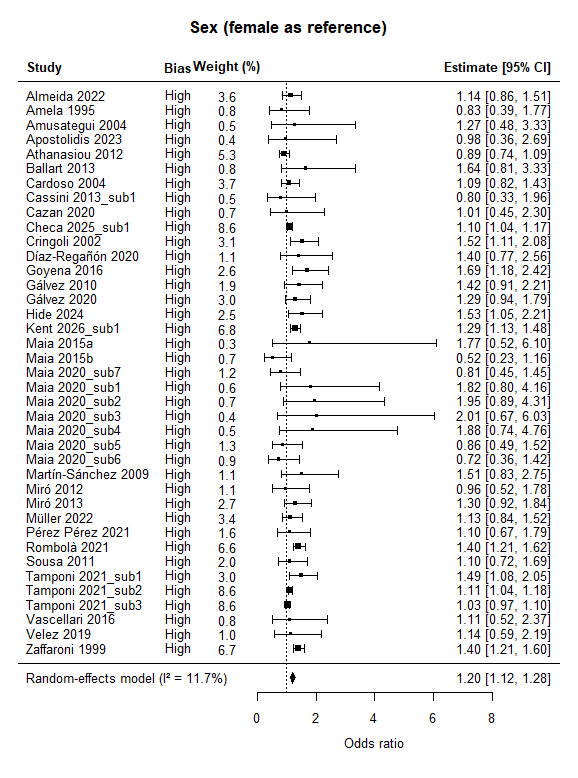

**Figure S8.** Random-effects forest plot of dog sex as a risk factor for *Leishmania* infection in dogs, based on sub-study specific *unadjusted* odds ratios. Plot elements are defined in Figure S3.

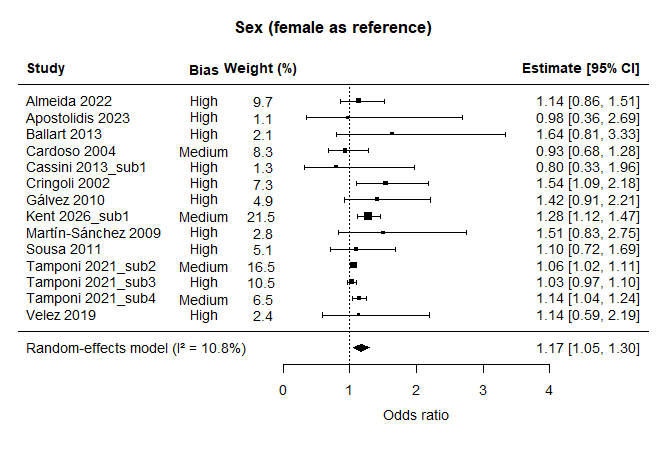

**Figure S9.** Random-effects forest plot of dog sex as a risk factor for *Leishmania* infection in dogs, based on sub-study specific *mixed-estimates* odds ratios. Plot elements are defined in Figure S3.

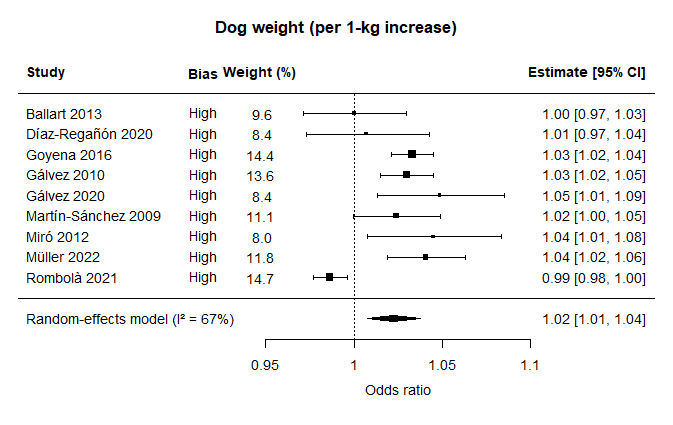

**Figure S10.** Random-effects forest plot of dog weight (per kg) as a risk factor for *Leishmania* infection in dogs, based on sub-study specific *unadjusted* odds ratios. Plot elements are defined in Figure S3.

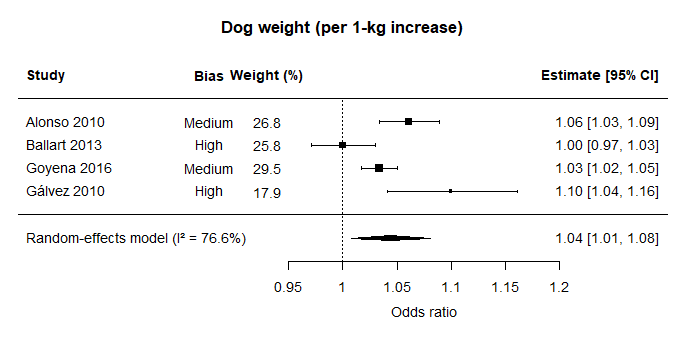

**Figure S11.** Random-effects forest plot of dog weight (per kg) as a risk factor for *Leishmania* infection in dogs, based on sub-study specific *mixed-estimates* odds ratios. Plot elements are defined in Figure S3.

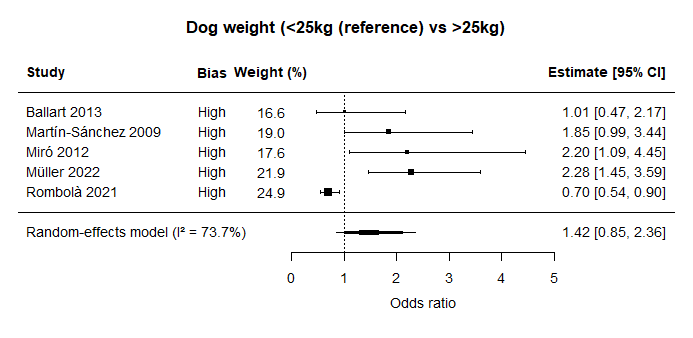

**Figure S12.** Random-effects forest plot of dog weight (<25kg vs >25kg) as a risk factor for *Leishmania* infection in dogs, based on sub-study specific *unadjusted* odds ratios. Plot elements are defined in Figure S3.

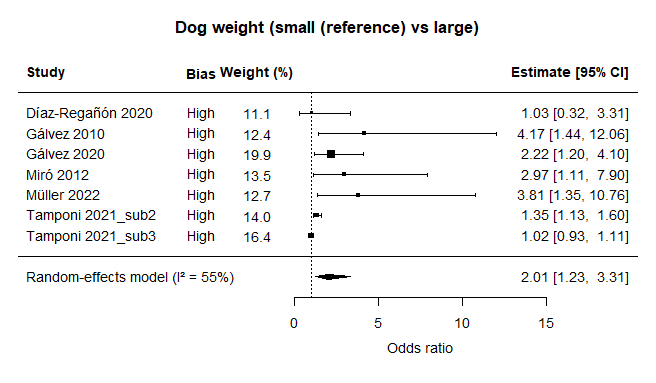

**Figure S13.** Random-effects forest plot of dog weight (small vs large) as a risk factor for *Leishmania* infection in dogs, based on sub-study specific *unadjusted* odds ratios. Plot elements are defined in Figure S3.

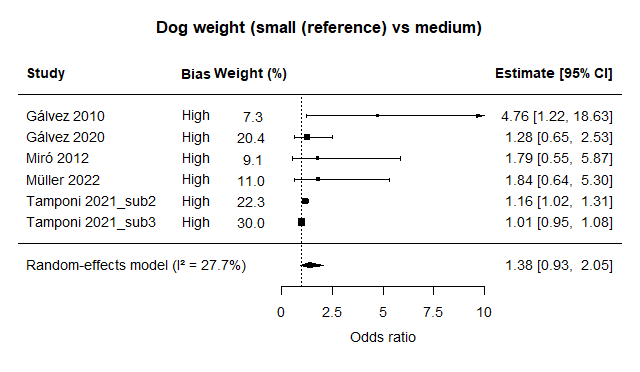

**Figure S14.** Random-effects forest plot of dog weight (small vs medium) as a risk factor for *Leishmania* infection in dogs, based on sub-study specific *unadjusted* odds ratios. Plot elements are defined in Figure S3.

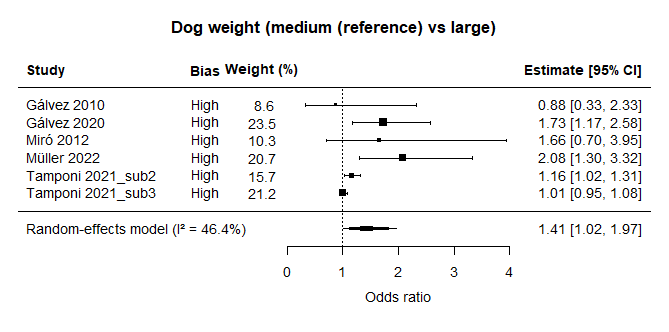

**Figure S15.** Random-effects forest plot of dog weight (medium vs large) as a risk factor for *Leishmania* infection in dogs, based on sub-study specific *unadjusted* odds ratios. Plot elements are defined in Figure S3.

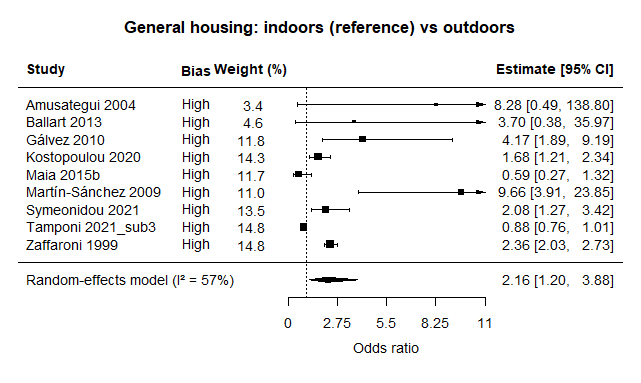
**Figure S16.** Random-effects forest plot of general housing (indoors vs outdoors) as a risk factor for *Leishmania* infection in dogs, based on sub-study specific *unadjusted* odds ratios. Plot elements are defined in Figure S3.

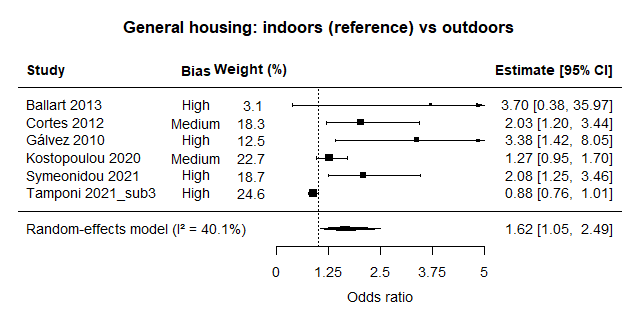
**Figure S17.** Random-effects forest plot of general housing (indoors vs outdoors) as a risk factor for *Leishmania* infection in dogs, based on sub-study specific *mixed-estimates* odds ratios. Plot elements are defined in Figure S3.

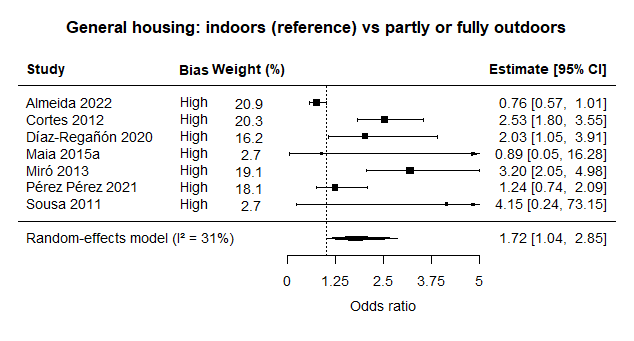
**Figure S18.** Random-effects forest plot of general housing (exclusively indoors vs partly or fully outdoors) as a risk factor for *Leishmania* infection in dogs, based on sub-study specific *unadjusted* odds ratios. Plot elements are defined in Figure S3.

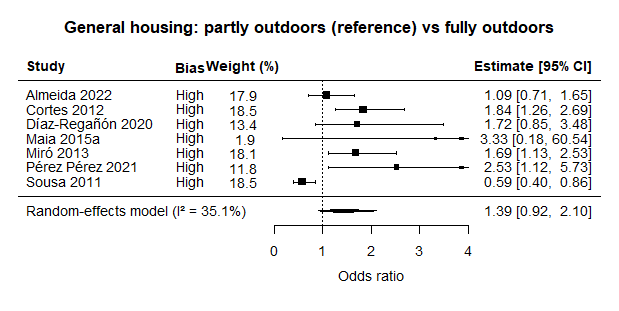
**Figure S19.** Random-effects forest plot of general housing (partly outdoors vs fully outdoors) as a risk factor for *Leishmania* infection in dogs, based on sub-study specific *unadjusted* odds ratios. Plot elements are defined in Figure S3.

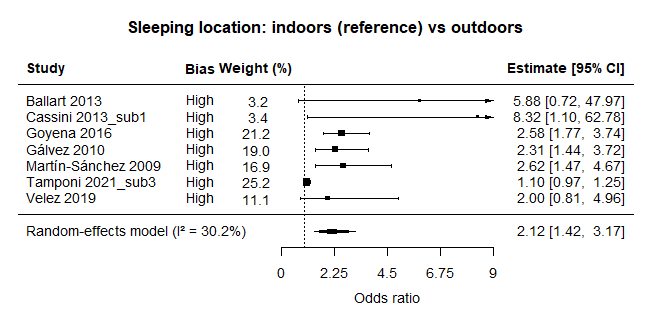

**Figure S20.** Random-effects forest plot of sleeping location (indoors vs outdoors) as a risk factor for *Leishmania* infection in dogs, based on sub-study specific *unadjusted* odds ratios. Plot elements are defined in Figure S3.

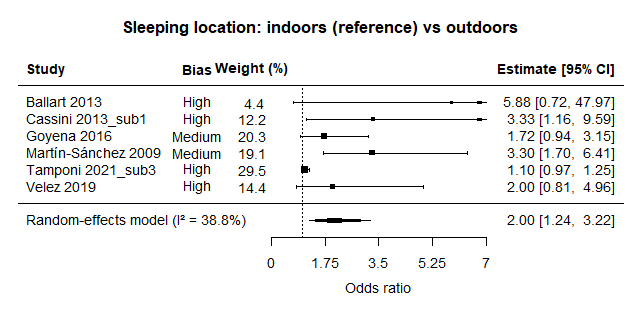

**Figure S21.** Random-effects forest plot of sleeping location (indoors vs outdoors) as a risk factor for *Leishmania* infection in dogs, based on sub-study specific *mixed-estimates* odds ratios. Plot elements are defined in Figure S3.

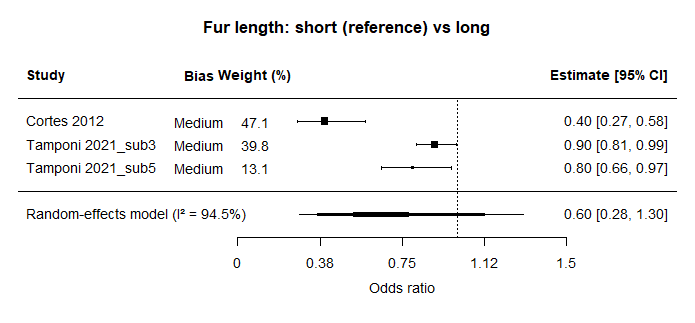

**Figure S22.** Random-effects forest plot of fur length (short vs long) as a risk factor for *Leishmania* infection in dogs, based on sub-study specific *adjusted* odds ratios. Plot elements are defined in Figure S3.

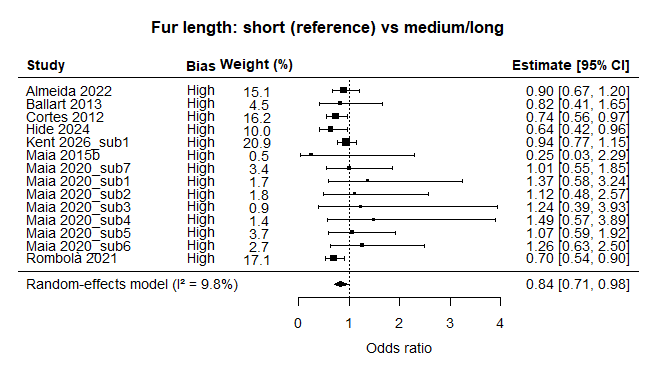

**Figure S23.** Random-effects forest plot of fur length (short vs medium/long) as a risk factor for *Leishmania* infection in dogs, based on sub-study specific *unadjusted* odds ratios. Plot elements are defined in Figure S3.

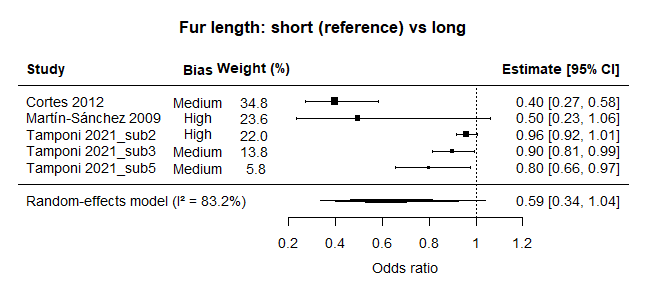

**Figure S24.** Random-effects forest plot of fur length (short vs long) as a risk factor for *Leishmania* infection in dogs, based on sub-study specific *mixed-estimates* odds ratios. Plot elements are defined in Figure S3.

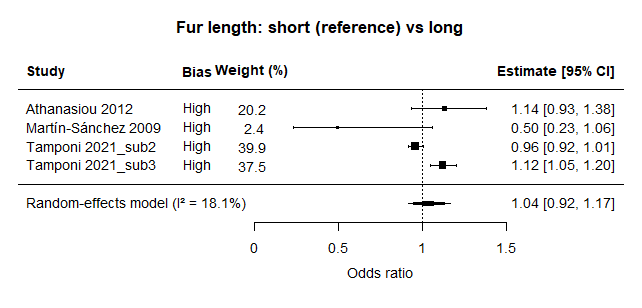

**Figure S25.** Random-effects forest plot of fur length (short vs long) as a risk factor for *Leishmania* infection in dogs, based on sub-study specific *unadjusted* odds ratios. Plot elements are defined in Figure S3.

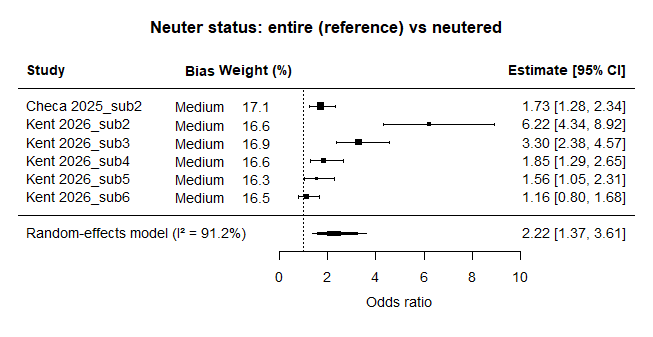

**Figure S26.** Random-effects forest plot of neuter status (entire vs neutered) as a risk factor for *Leishmania* infection in dogs, based on sub-study specific *adjusted* odds ratios. Plot elements are defined in Figure S3. Kent 2026 sub-studies are ordered after dog age, with sub2 representing 1-3 year olds, sub3 represents 3-5 years, sub4 represents 5-7 years, sub5 represents 7-9 years and sub6 represents 9 years or older.

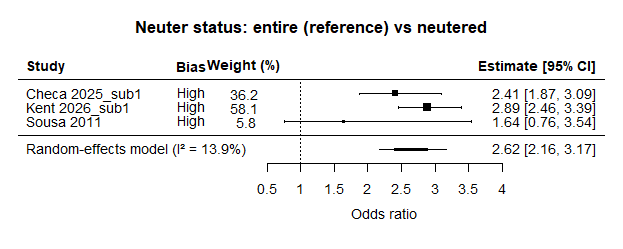

**Figure S27.** Random-effects forest plot of neuter status (entire vs neutered) as a risk factor for *Leishmania* infection in dogs, based on sub-study specific *unadjusted* odds ratios. Plot elements are defined in Figure S3.

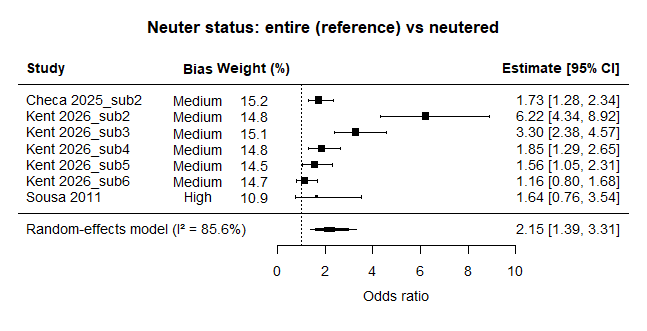

**Figure S28.** Random-effects forest plot of neuter status (entire vs neutered) as a risk factor for *Leishmania* infection in dogs, based on sub-study specific *mixed-estimates* odds ratios. Plot elements are defined in Figure S3.

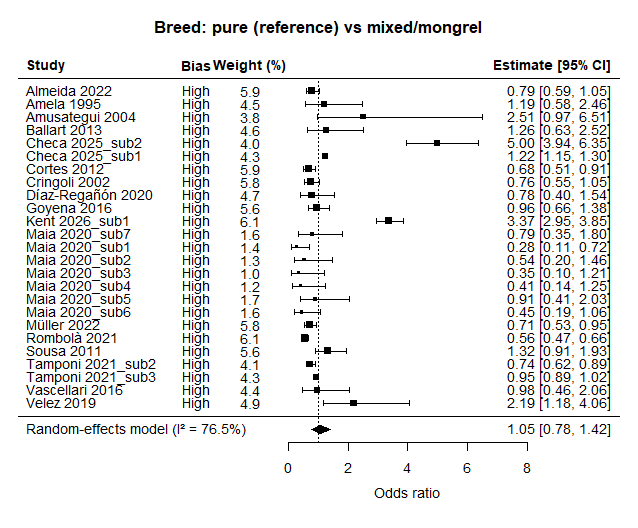

**Figure S29.** Random-effects forest plot of breed (pure vs mixed/mongrel) as a risk factor for *Leishmania* infection in dogs, based on sub-study specific *unadjusted* odds ratios. Plot elements are defined in Figure S3.

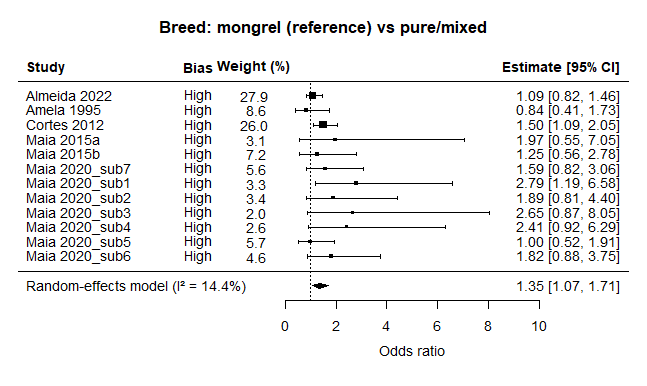

**Figure S30.** Random-effects forest plot of breed (mongrel vs pure/mixed) as a risk factor for *Leishmania* infection in dogs, based on sub-study specific *unadjusted* odds ratios. Plot elements are defined in Figure S3.

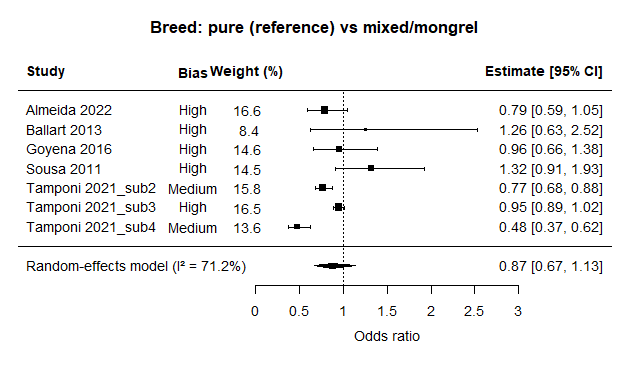

**Figure S31.** Random-effects forest plot of breed (pure vs mixed/mongrel) as a risk factor for *Leishmania* infection in dogs, based on sub-study specific *mixed-effects* odds ratios. Plot elements are defined in Figure S3.

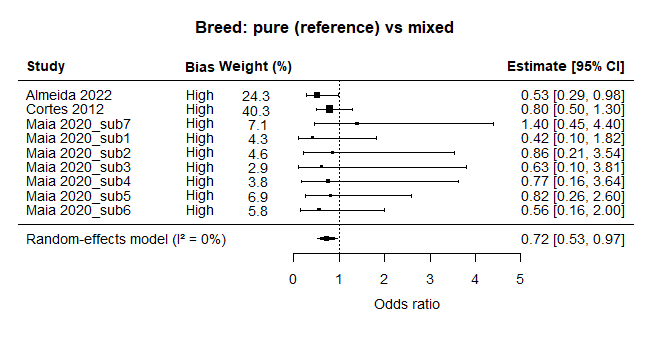

**Figure S32.** Random-effects forest plot of breed (pure vs mixed) as a risk factor for *Leishmania* infection in dogs, based on sub-study specific *unadjusted* odds ratios. Plot elements are defined in Figure S3.

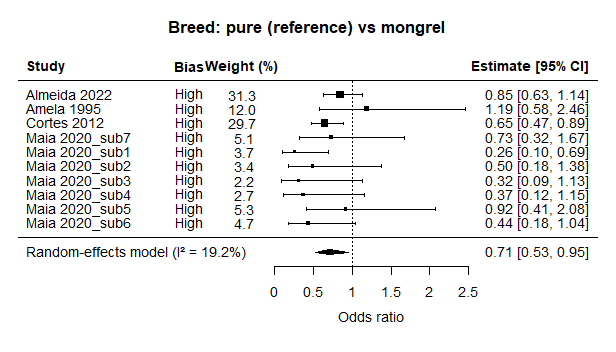

**Figure S33.** Random-effects forest plot of breed (pure vs mongrel) as a risk factor for *Leishmania* infection in dogs, based on sub-study specific *unadjusted* odds ratios. Plot elements are defined in Figure S3.

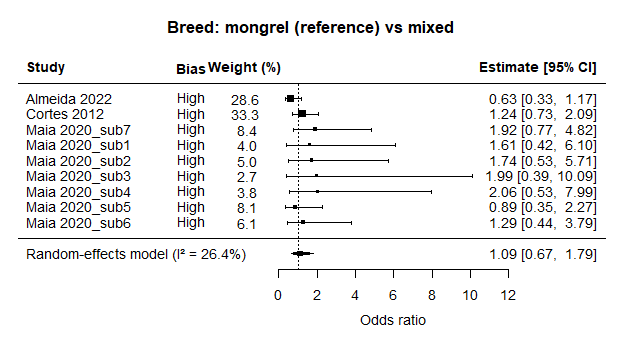

**Figure S34.** Random-effects forest plot of breed (mongrel vs mixed) as a risk factor for *Leishmania* infection in dogs, based on sub-study specific *unadjusted* odds ratios. Plot elements are defined in Figure S3.

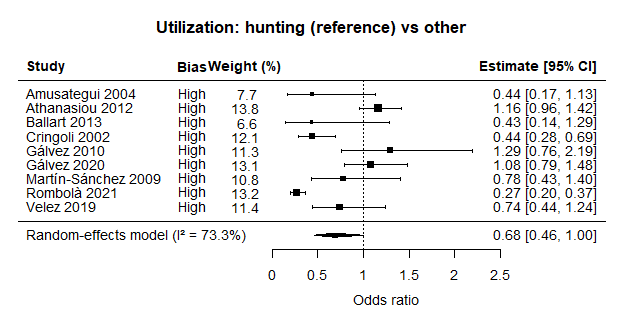
**Figure S35.** Random-effects forest plot of utilization (hunting vs other) as a risk factor for *Leishmania* infection in dogs, based on sub-study specific *unadjusted* odds ratios. Plot elements are defined in Figure S3.

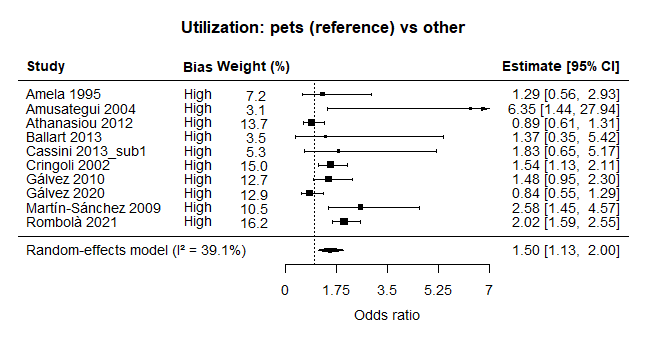
**Figure S36.** Random-effects forest plot of utilization (pets vs other) as a risk factor for *Leishmania* infection in dogs, based on sub-study specific *unadjusted* odds ratios. Plot elements are defined in Figure S3.

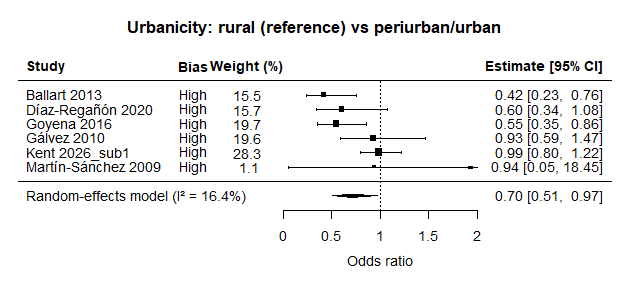

**Figure S37.** Random-effects forest plot of urbanicity (rural vs periurban/urban) as a risk factor for *Leishmania* infection in dogs, based on sub-study specific *unadjusted* odds ratios. Plot elements are defined in Figure S3.

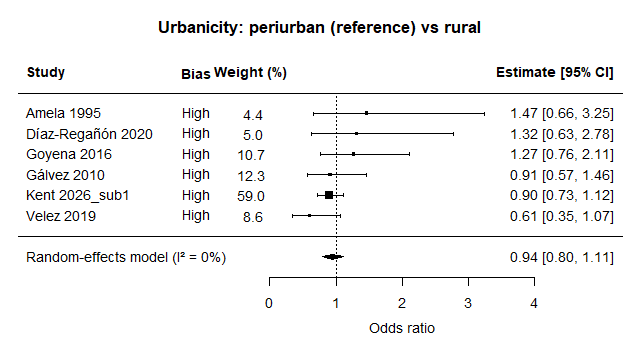

**Figure S38.** Random-effects forest plot of urbanicity (periurban vs rural) as a risk factor for *Leishmania* infection in dogs, based on sub-study specific *unadjusted* odds ratios. Plot elements are defined in Figure S3.

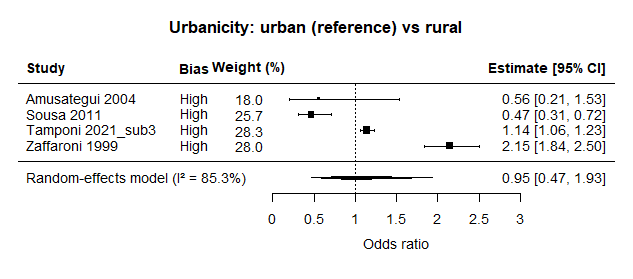

**Figure S39.** Random-effects forest plot of urbanicity (urban vs rural) as a risk factor for *Leishmania* infection in dogs, based on sub-study specific *unadjusted* odds ratios. Plot elements are defined in Figure S3.

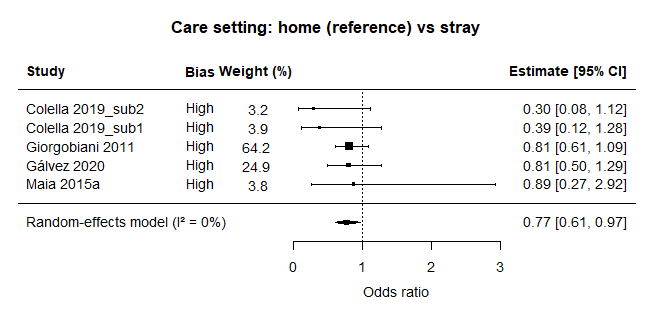
**Figure S40.** Random-effects forest plot of care setting (home vs stray) as a risk factor for *Leishmania* infection in dogs, based on sub-study specific *unadjusted* odds ratios. Plot elements are defined in Figure S3.

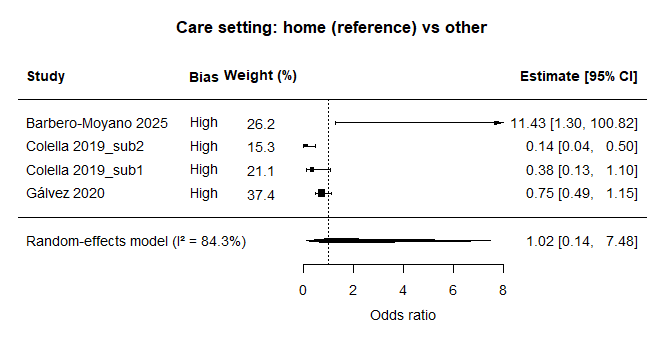
**Figure S41.** Random-effects forest plot of care setting (home vs other) as a risk factor for *Leishmania* infection in dogs, based on sub-study specific *unadjusted* odds ratios. Plot elements are defined in Figure S3.

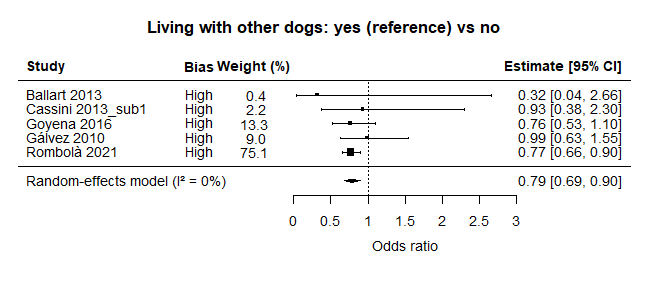

**Figure S42.** Random-effects forest plot of cohabitation with other dogs (yes vs no) as a risk factor for *Leishmania* infection in dogs, based on sub-study specific *unadjusted* odds ratios. Plot elements are defined in Figure S3.

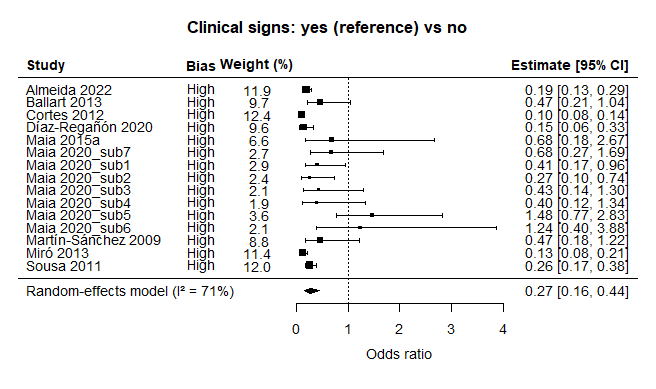

**Figure S43.** Random-effects forest plot of clinicals signs (yes vs no) as a risk factor for *Leishmania* infection in dogs, based on sub-study specific *unadjusted* odds ratios. Plot elements are defined in Figure S3.

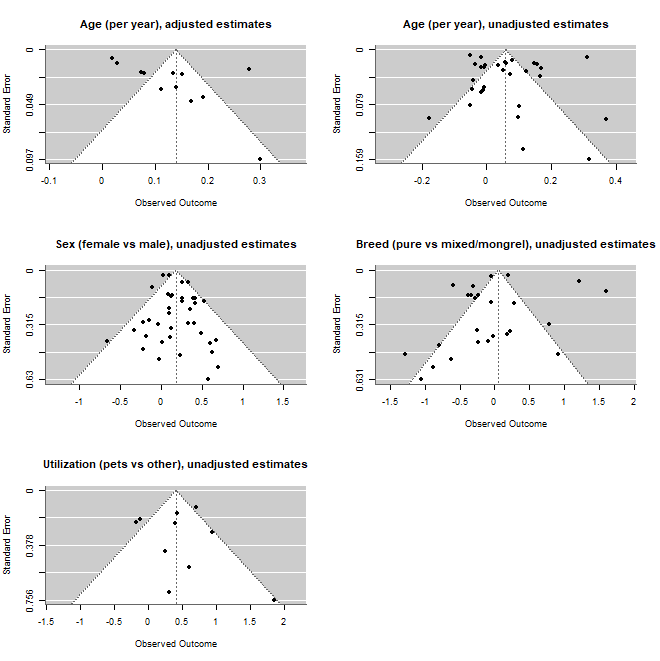

**Figure S44.** Funnel plots for risk factors with estimates from at least 10 studies. Each panel corresponds to an analytical unit where a random-effects model was used. Log odds ratio (OR) on the x-axes, and standard error of log OR on the y-axes. Each point represents a sub-study.
